# A single valine to leucine switch disrupts *Plasmodium falciparum* AP2-G DNA binding and reveals GDV1’s role in *ap2-g* activation

**DOI:** 10.1101/2025.04.22.648970

**Authors:** Surendra K. Prajapati, Jeffrey X. Dong, Belinda J. Morahan, Thoai Dotrang, Michelle C. Barbeau, April E. Williams, Daniel Hupalo, Matthew D. Wilkerson, Clifton L. Dalgard, Bjorn F.C. Kafsack, Manuel Llinás, Kim C. Williamson

## Abstract

Sexual commitment in *Plasmodium* parasites is essential for malaria transmission, yet much remains unknown about the underlying signaling events initiating sexual conversion in a subpopulation of parasites. We discovered a single valine (V_2163_) to leucine (L_2163_) mutation in an Apetala 2 (AP2) transcription factor required for *P. falciparum* gametocytogenesis, *ap2-g* that abrogates sexual differentiation and confirmed this with forward and reverse mutation editing. Mutated AP2-G.L_2163_ does not bind the *ap2-g* consensus motif, GnGTAC, or stimulate AP2-G-dependent gene transcription including autoregulation. We then used AP2-G.L_2163_ parasite lines as tools to demonstrate the critical role of GDV1 in the initial activation of the silent *ap2-g* locus during the trophozoite to schizont transition in the absence of functional AP2-G and its autoregulation. Additionally, we show that AP2-G.V is required for MSRP1 expression, which can be used to distinguish early and late sexually committed schizonts. Together this work demonstrates that valine_2163_ in AP2-G plays a critical role in DNA binding, highlighting the functional importance of this specific region for malaria transmission as well as the critical role of GDV1 in the initial activation of *ap2-g* expression and induction of sexual differentiation. The reporter lines generated allow further study of signaling pathways or screening of factors regulating sexual commitment.

## Introduction

Malaria transmission is a global public health concern which accounted for 249 million clinical cases and more than half a million of deaths in 2022 ^1^. A critical step in human to human transmission via mosquito is the production of sexual stage *Plasmodium* parasites, called gametocytes. Sexual differentiation begins in a subpopulation of blood stage parasites during the asexual replication cycle, but the percent of parasites that commit to sexual development varies in vitro and in vivo^2, 3^. In contrast to relatively low in vitro gametocyte-commitment rates (GCR) (<30%)^2^ in *P. falciparum* laboratory adapted strains and clinical isolates, in vivo symptomatic or asymptomatic malaria studies show dramatic variability in GCRs (0-78%)^4, 3^. The factors underlying this quantitative variability in gametocyte production remain undefined and make it difficult to target and eliminate the infectious reservoir. This GCR variation coupled, with the high prevalence of asymptomatic malaria infections, which can be as high as 60-80% in some regions^5, 6, 7^, and the inability of commonly used antimalarials to kill mature gametocytes^8^ are major challenges to malaria control programs. Defining the molecular mechanisms underlying the regulation of sexual commitment should facilitate the design of new transmission blocking strategies.

Two genes have been found to be required to initiate sexual commitment in *P. falciparum*, *gametocyte development protein 1* (*gdv1*: PF3D7_0935400) ^9^ and a plant-like *apetala 2* transcription factor (*ap2-g*: PF3D7_1222600) ^10^. GDV1 and AP2-G are required for *ap2-g* expression ^3, 11^, while AP2-G also activates downstream genes needed for the progression of sexually committed rings to continue gametocyte development ^10, 12^. In asexually-committed blood stage parasites, the *ap2-g* locus is epigenetically silenced by histone 3 lysine 9 trimethylation (H3K9Me3)^13^ that is stabilized by heterochromatin protein 1 (HP1: PF3D7_1220900) ^11^ ^14^. As the blood stage parasite matures, the start of DNA replication marks entry into the schizont stage of the life cycle. During schizogony, it has been proposed that in the subpopulation of schizonts that express GDV1, the *ap2-g* locus is de-repressed ^11^, but the induction and regulation of this decision has not been established. There is evidence that once GDV1 is expressed, it interacts with HP-1 and this interaction has been proposed to contribute to *ap2-g* de-repression by removing HP1 ^11^, but the detailed mechanism that confers specificity to the release of HP-1 only at the *ap2-g* locus and only in some parasites remains to be elucidated. Once expressed AP2-G induces its own expression, termed autoregulation, by binding to the core AP2-G DNA recognition motif GnGTAC that is present in the 5’ flanking region of *ap2-g* ^10, 12^. This motif is also present upstream of a number of other genes, some of which are upregulated in the presence of AP2-G in schizonts and/or the next generation of blood stage parasites, called rings, initiating gametocyte differentiation. The reasons different AP2-G-dependent genes are expressed at different times after AP2-G expression remain unclear as does the regulation of initial expression of GDV1 and AP2-G.

In higher eukaryotes, de-repression of epigenetically silent genes has been described through post-translation modifications ^15^. Phosphorylation of histone 3 serine 10 has been demonstrated to play a critical role in release of HP1 from lysine 9 trimethylated H3 ^16^ ^17, 18^. This change allows RNA polymerase access to transcribe the previously silenced genes. In *P. falciparum,* several studies have consistently shown phosphorylation of H3 serine-29 and 33 ^19, 20, 21^ in trophozoite and schizont stages, which is when sexual commitment is also thought to be initiated. It is possible that phosphorylation of H3 serine 29 and/or 33 or both could play critical roles in releasing HP1 from *ap2-g* locus bound H3K9Me3.

We sought to evaluate whether serine/threonine kinase (STK2: PF3D7_0214600), which has a nuclear localization signal, could be one of the kinases playing a role in the phosphorylation of H3 ^20, 22^. We used reverse genetics to define the role of STK2 in gametocytogenesis but, unexpectedly, whole genome sequencing of the *stk2* knockout parasites generated revealed a single valine_2163_ to leucine_2163_ mutation in *ap2-g*. We show that the replacement of valine with leucine in the presence and absence of *skt2* completely abrogates AP2-G binding to the upstream core GnGTAC motif, autoregulation, downstream gene expression and gametocyte production. The development of additional mutant and wildtype *ap2-g* reporter lines with and without inducible GDV1 expression allowed quantification of the initiation of *ap2-g* expression in the absence of functional AP2-G and its autoregulation, emphasizing the critical role of GDV1 in the induction of sexual commitment.

## Results

### Determining the genetic locus underlying a complete loss of gametocytogenesis

In the course of evaluating kinases for a role in gametocytogenesis, we generated a serine/threonine kinase (STK2: PF3D7_0214600) knockout line (*3D7.stk2Δ*) in a gametocyte-competent *P. falciparum* 3D7 strain **(Fig S1a-c).** Asexual growth was normal consistent with the previous piggyBac transposon insertion mutagenesis prediction that *stk2* is dispensable during asexual growth ^23^. In marked contrast, *3D7.stk2Δ* clones did not produce immature or mature gametocytes (**Fig 1a-b, Fig S1d**). To determine whether the block was at sexual commitment or maturation of sexually committed rings, we quantified gametocyte-committed ring stage parasites using the validated early gametocyte biomarker *ap2-g* ^4^. *Ap2-g* expression levels were significantly reduced in *3D7.stk2Δ* ring stage parasites (**Fig 1c**), suggesting a block in sexual commitment similar to that observed in previously reported gametocyte deficient lines, F12 which has a premature stop codon in *ap2-g* at bp 3923^10^, and a gametocyte inducible line *NF54.gdv1.3xha.fkbp* in the absence of Shield1 (Shld1-). To confirm the phenotype and define the cellular localization of STK2, we used the original double crossover plasmid to generate an inducible STK2 reporter line (*NF54.stk2.gfp.fkbp*) as well as a standard knock out (*NF54.stk2Δ*) in a gametocyte competent Pf strain NF54. Surprisingly, the *NF54.stk2Δ* and *NF54.stk2.gfp.fkbp* lines with and without Shld1 all produced gametocytes and had GCRs similar to the parental line NF54 (**Fig 1d-e**). These results strongly suggest that the loss of gametocytogenesis in *3D7.stk2Δ* clones was independent of *stk2*.

**Fig. 1:**
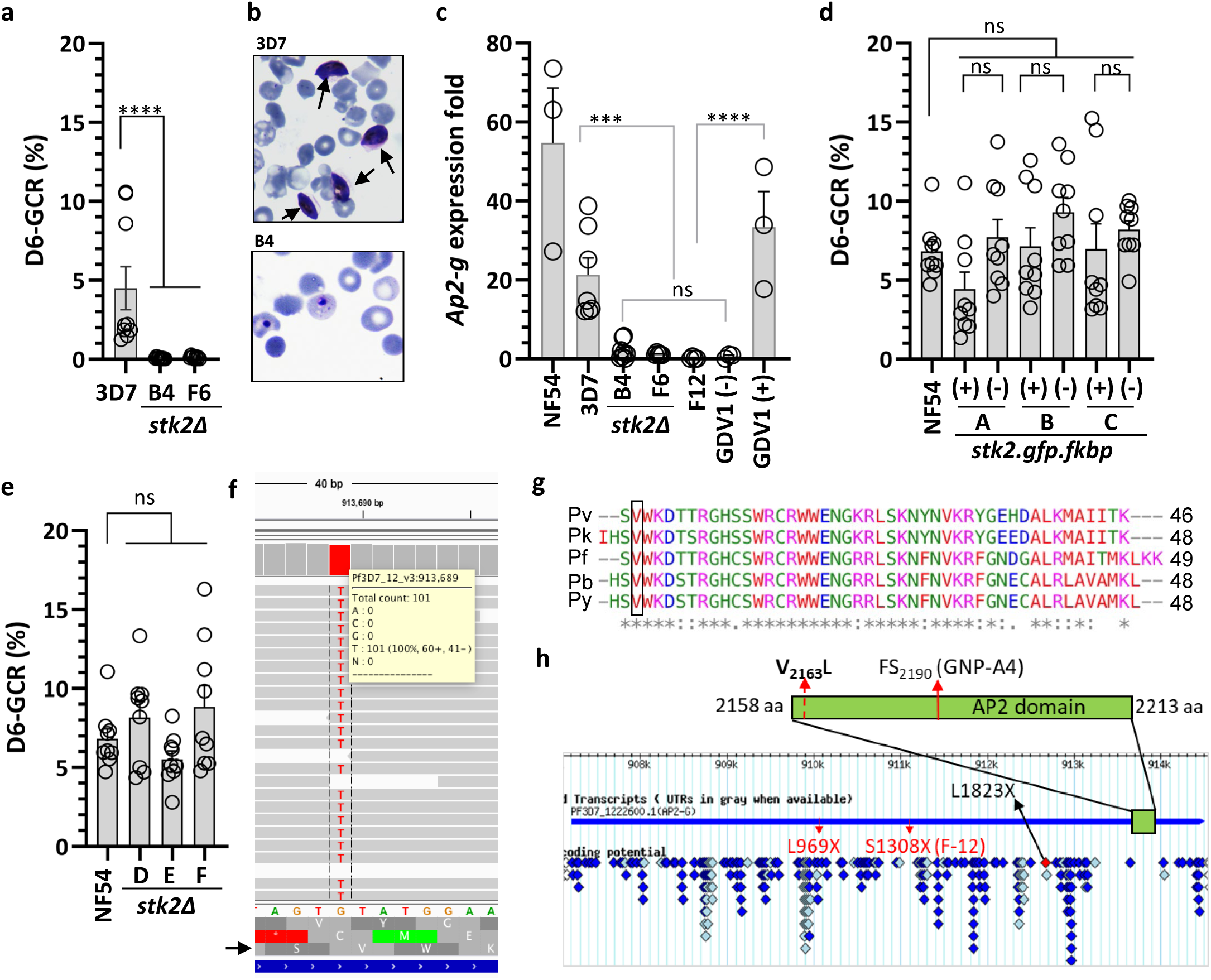
Role of Serine/Threonine protein kinase (*Pfstk2)* in gametocytogenesis. **a)** Gametocyte conversion rate (GCR) of *3D7.stk2Δ* clones (B4, F6) and the parental line (3D7). **b)** NAG-D6 thin smear showing the presence/absence of gametocytes (arrows) in the parent 3D7 and *3D7.stk2Δ* clone (B4). **c)** *Ap2-g* expression fold changes in ring stage parasites from NF54, 3D7, *3D7.stk2Δ* clones (B4, F6), *ap2-g* truncated line F12, and *NF54.gdv1.3xha.fkbp* in the absence (GDV1-) or presence (GDV1+) of Shld1. **d)** GCR phenotype of the parent NF54 and STK2 inducible *NF54.stk2.gfp.fkbp* clones (A, B & C) in the presence (+) or absence (-) of Shld1. **e)** GCR phenotype of the parent NF54 and *NF54.stk2Δ* clones (D, E & F). **f)** *Ap2-g* mutation (g.6487G>T) resulting in p.V2163L amino acid substitution was present in all the sequence reads from the *3D7.stk2Δ* line (gray bars). The 3D7 reference sequence is shown at the bottom just above the corresponding amino acid sequence in all 3 reading frames. The arrow indicates the third reading frame encodes AP2-G. **g)** Protein sequence alignment of the AP2-G AP2 domain derived from five diverse malaria parasite species, *P. vivax* (Pv), *P. knowlesi* (Pk), *P. falciparum* (Pf), *P. berghei* (Pb) and *P. yoellii* (Py), indicates conservation of the mutated valine (black box). **h)** Schematic of AP2-G (aa 1 - 2432) with the AP2 DNA binding domain indicated by a green box and SNPs previously identified in the *ap2-g* locus in lab-adapted lines (red arrow), the present study (red dotted arrow) or field isolates (diamonds ^22^). For graphs a, c, d and e independent data points are indicated by circles and the bar and error bar represent the mean and standard error of the mean, respectively. A Kruskal-Wallis test followed by Dunn’s multiple comparison was used for statistical analysis between groups. The p value of each comparison is indicated by ns (>0.05), * (<0.05), *** (<0.001), and **** (<0.0001).

The *gdv1* and *ap2-g* loci previously associated with gametocyte production ^9, 10^, remained intact in the *3D7.stk2Δ* line and were free from the known *ap2-g* nonsense (F12) or frameshift (GNP-4) mutations ^10^, or nonsense mutations (K561*, Q578*) in *gdv1* ^24, 25^ (**Fig S2**). To extend our search, we sequenced the whole genome of the *3D7.stk2Δ* line. Whole genome analysis confirmed the lack of deletions or mutations conferring premature termination of *gdv1* or *ap2-g*. However, *ap2-g* had a missense mutation (g.6487G>T) (**Fig 1f**) that resulted in a switch from valine (V) to leucine (L) in the AP2 domain (V2163L). This V_2163_L mutation was the only missense SNP identified in *3D7.stk2Δ* not present in wild type NF54 (**Table S1**). The AP2-G AP2 domain is 50 aa long (AP2-G_2162-2210_) and highly conserved across the *Plasmodium* species infecting human, rodent and monkeys with all these species having a valine at the position corresponding to V_2163_ (**Fig 1g**). Additionally, in the global *P. falciparum* isolate repositories analyzed to date, no mutations have been reported in the AP2-G AP2 domain (PlasmoDB^22^, MalariaGEN^26^) (**Fig 1h**). These results suggest that the AP2-G AP2 domain is under a strong selective pressure in the field as expected for a gene required for transmission. The *3D7.stk2Δ* line was therefore designated as *3D7.stk2Δ//ap2-g.L* and wild type (wt) or mutant *ap2-g* designated as *ap2-g.V* or *ap2-g.L*, respectively.

### The AP2-G V_2163_L substitution is sufficient for a loss of gametocytogenesis

To confirm that *ap2-g.V_2163_L* prevents gametocytogenesis, we swapped the V allele with an L allele in a gametocyte competent *P. falciparum* strain NF54 using a forward mutation editing approach (**Fig 2a**), generating transgenic mutant lines (*NF54.ap2-g.L, NF54.ap2-g.L.gfp.fkbp*). While both lines have a V to L replacement, the *NF54.ap2-g.L.gfp.fkbp* line also has GFP.FKBP inserted in frame to regulate AP2-G.L. The whole genomes of both lines were sequenced and compared to the parental line, NF54, to rule out the presence of any unintended mutations. None of the lines expressing AP2-G.L produced immature or mature gametocytes (**Fig 2b-c**) which is the same phenotype shown by original *3D7.stk2Δ//ap2-g.L* clones (**Fig 1a-b**). To strengthen our hypothesis, we also tested whether reversing the L_2163_ mutation back to V can restore gametocytogenesis in the *3D7.stk2Δ//ap2-g.L* line, as well as the *NF54.ap2-g.L.gfp.fkbp* line **(Fig 2d)**. Correcting the mutation to V restored the rate of gametocytogenesis to that observed in the original parental lines (NF54 or 3D7) (**Fig 2e-f**). This forward and reverse mutation editing approach firmly supports the valine to leucine mutation, not the *stk2* knockout or another mutation, as the cause of the loss of gametocytogenesis in the *3D7.stk2Δ//ap2-g.L* line.

**Fig 2:**
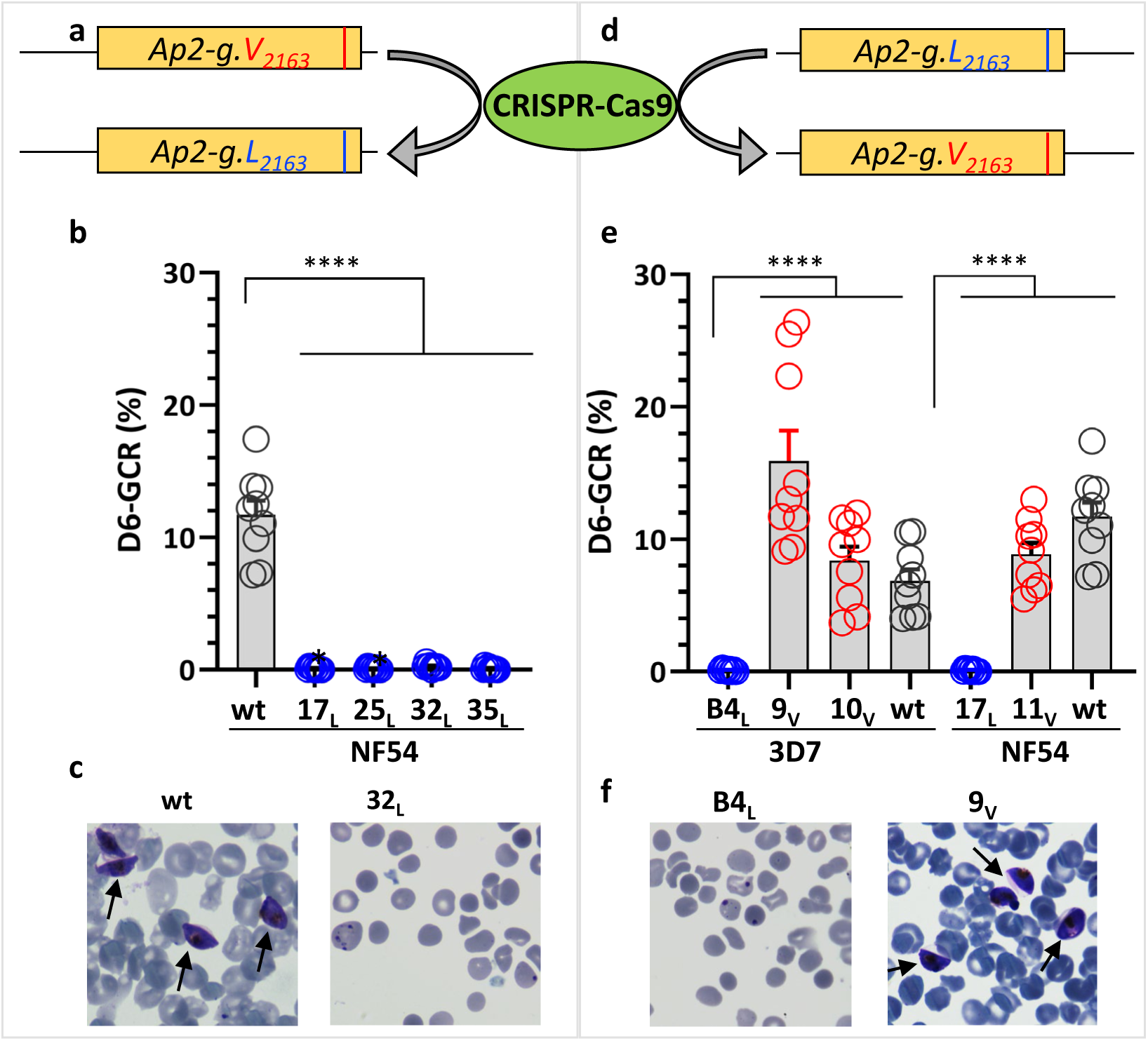
Forward and reverse editing confirms AP2-G.L causes loss of sexual conversion. **a)** Schematic of the allele swapping strategy showing the exchange of wild type (wt) *ap2-g* allele V with the mutant allele (L) in strain NF54. **b)** Day 6 (D6) gametocyte conversion rate (GCR) analysis of wt and mutated NF54 parasites, *NF54.ap2-g.L* clones 32_L_& 35_L_ were generated using CRISPR-Cas9 and *NF54.ap2-g.L.gfp.fkbp:* clones 17_L_ & 25_L_ were generated using homologous recombination without CRISPR-Cas9 and grown in the presence of Shld1. **c)** D6 Giemsa-stained smear of wt NF54 or mutated *NF54.ap2-g.L* clone 32_L_. **d)** Schematic of allele swapping strategy for the reversion of mutant allele L into wild type V. **e)** Day 6 GCR analysis of 3D7 or NF54 derived lines with wt (V) or mutated (L) *ap2-g* lines, *3D7.stk2Δ//ap2-g.L* clone B4_L_, *3D7.stk2Δ//ap2-g.V* clones 9_V_ & 10_V_ and *NF54.ap2-g.L.gfp.fkbp* clone 17_L_ and *NF54.ap2-g.V* clone 11_V_. **f)** D6 Giemsa stained smear of 3D7, L mutated line (*3D7.stk2Δ//ap2-g.L* clone B4_L_), and mutation corrected line (*3D7.stk2Δ//ap2-g.V* clone 9_V_). Gametocytes are indicated by an arrow. Blue, red, or black circles indicate mutated (*Ap2-g.L*), mutation corrected (*Ap2-g.V*), or parent with wt *ap2-g* lines, respectively. Each circle indicates an independent data point and the bar and error bar represent the mean and standard error of the mean, respectively. A Kruskal-Wallis test followed by Dunn’s multiple comparison was used for statistical analysis between groups. The p value of each comparison is indicated by ns (>0.05) and **** (<0.0001).

### V_2163_L switch disrupts DNA binding

In silico analysis indicates V_2163_ is the second aa in the predicted DNA binding region located at the N terminus of the full AP2 domain (aa 2162 to 2210) (**Fig 3a-b**). Changing V_2163_ to L to mimic the observed mutation shifted the start of the predicted AP2 domain downstream by 11 aa which decreases size and interaction strength (i-Evalue) of the predicted DNA binding region (**Fig 3b**). Eleven aa shorter AP2 domains were also predicted when V_2163_ was substituted by any of the other aa (**Fig 3b, Fig S3**), except isoleucine which was predicted to cause only a six aa shift (**Fig 3b**). These computational predictions suggest that V_2163_ could be critical for DNA binding. To confirm the in-silico prediction, we generated recombinant AP2 domain protein (AP2-G_2150-2220_, 60 aa) containing mutated (AP2-G.L) or wild type amino acid (AP2-G.V) as well as *P. berghei* wild type PbAP2-G. The recombinant proteins were tested in an electrophoretic mobility shift assay (EMSA) for binding to a biotinylated oligonucleotide (GTGTACAC) that includes the core PfAP2-G DNA recognition motif GnGTAC ^12^ ^10^. In contrast to the wild type recombinant PfAP2 domain (AP2-G.V) and PbAP2 domain, the mutant recombinant protein (PfAP2-G.L) did not bind the GTGTACAC oligo (**Fig 3c**). Both the in vitro and in silico results suggest that mutated AP2-G.L_2163_ would not bind to this sequence in genomic DNA and therefore no longer act as a functional transcription factor to activate gene expression by binding to the core motif GnGTAC. The lack of DNA binding is also consistent with the inability of AP2-G.L to support *ap2-g* transcription and gametocyte production in the *3D7.stk2Δ//ap2-g.L* line **(Fig 1a-c)**.

**Fig 3:**
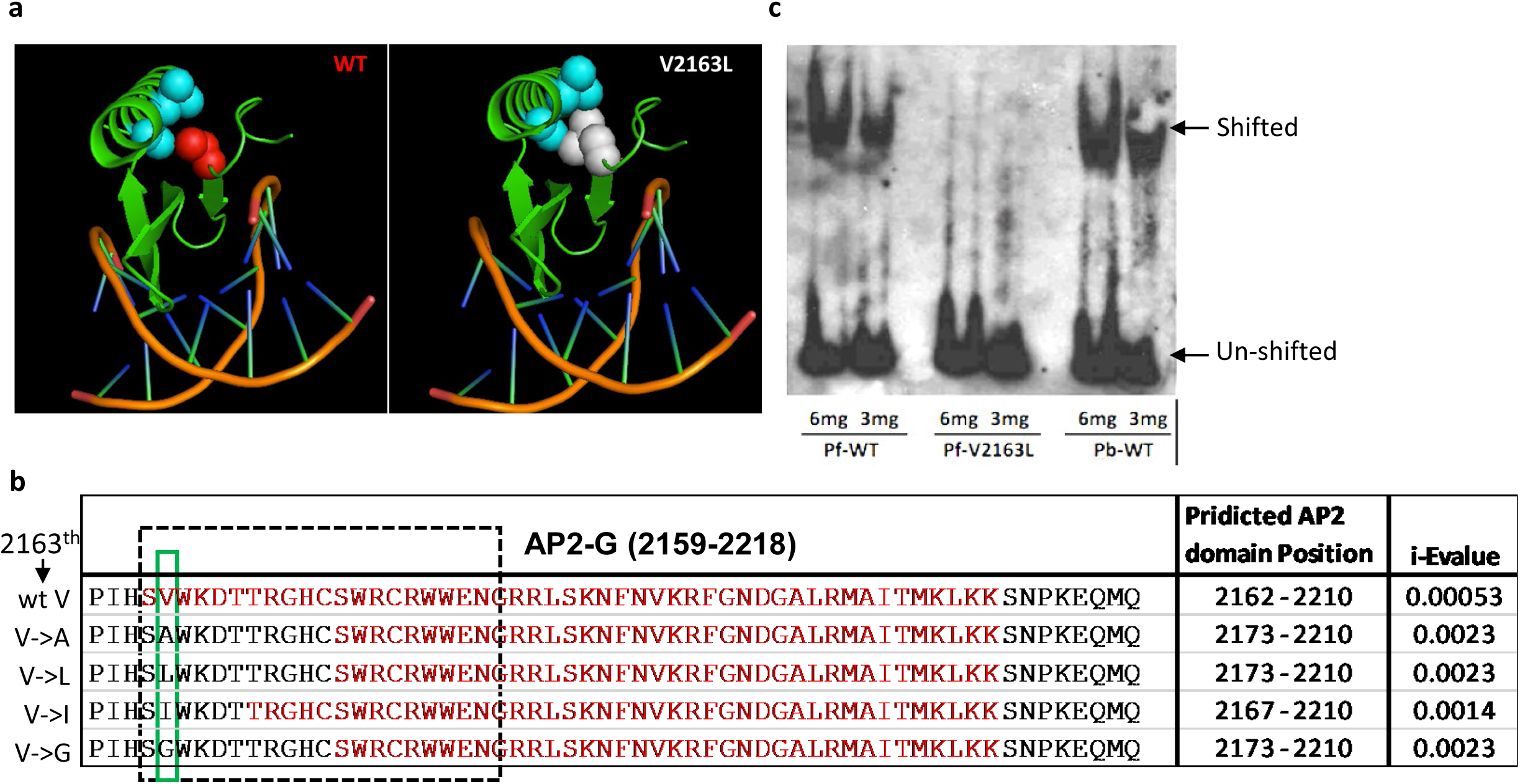
AP2-G.L does not bind DNA. **a)** In silico visualization of either the wild type valine or mutant leucine amino acid at the interface of AP2-G-DNA binding complex. **b)** Computational prediction of the amino acids (aa) contributing to the AP2 domain (red text), the DNA binding domain (dotted box), and the binding strength (i-Evalue) of the wild type *ap2-g* region (wt V) or following mutation of the indicated aa at position 2163 (green box). **c)** Gel shift analysis using a light shift EMSA assay kit with biotinylated probe GTGTACAC and recombinant proteins with either a wild type or mutated AP2 domain, *P. falciparum* recombinant AP2 domain, aa 2150-2220 (Pf-WT or Pf-V2163L) or *P. berghei* wild type AP2 domain (Pb-WT) ^59^. Shifted and unshifted bands are indicated by respective arrow.

### GDV1 activates *ap2-g* in the absence of AP2-G or its autoregulation

Parasite lines expressing AP2-G.L provide tools to investigate the initiation of *ap2-g* expression in the absence of AP2-G autoregulation and interrogate the separate roles of GDV1 and AP2-G in this initiation process. *Ap2-g* expression is highly dynamic during schizogony, therefore first we did an RNA time-course using a gametocyte inducible line (*NF54.gdv1.3xha.fkbp//ap2-g.V*) expressing wild type AP2-G to identify the best time to compare expression between lines. Prior to the experiments, parasites were maintained in the absence of Shld1 to eliminate contamination from gametocyte-committed parasites produced during earlier asexual cycles that also express *ap2-g* and could interfere in accurate determination of *ap2-g* expression initiation. After Shld1 addition to half the parasite cultures, *ap2-g* transcription was monitored every other hour starting from 28 hours post invasion (hpi) through 42 hpi in both Shld1 (+) and (-) conditions (**Fig 4a**). This time course covers the trophozoite to schizont developmental transition. The *ap2-g* RNA time-course revealed that in the presence of Shld1-stabilized GDV1, *ap2-g* transcripts first increased over baseline (∼two-fold) at 32 hpi and this increase was maintained until 36 hpi. After 36 hpi, *ap2-g* transcripts rose dramatically, until plateauing at 40 hpi. Six days later stage III-IV gametocytes were detected in the culture **(Fig 4b).** In contrast, in the absence of Shld1 there was no increase in *ap2-g* transcript (**Fig 4a**) and gametocytes were not produced (**Fig 4b**).

**Fig 4:**
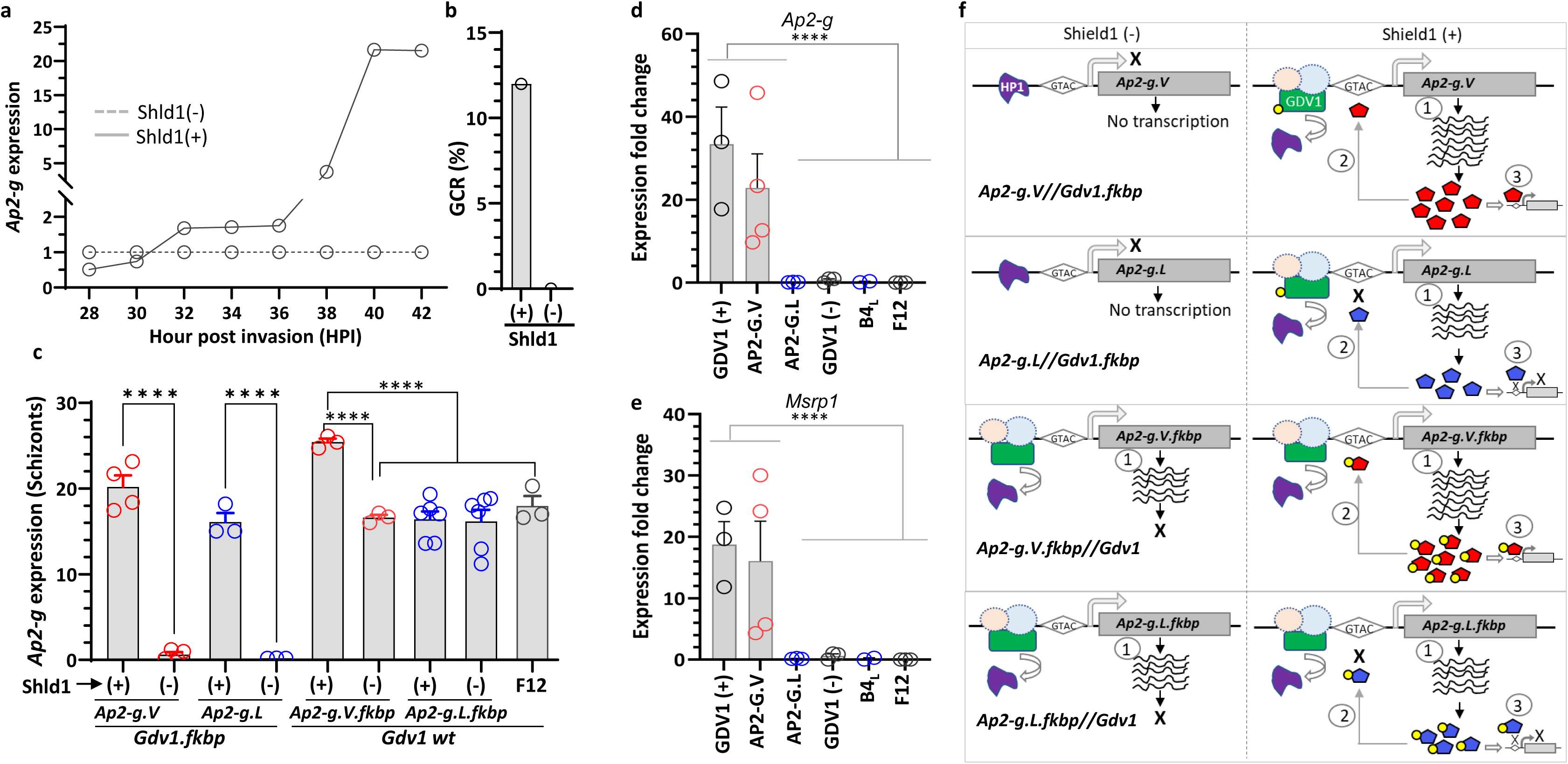
Engineered inducible *gdv1* and *ap2-g* lines demonstrate that both *ap2-g.V* and *.L* expression is GDV1 regulated. **a)** A time-course using the *NF54.gdv1.3xha.fkbp* line in absence and presence of Shld1 showing *ap2-g* transcript analysis every 2 hours starting from 28 to 42 hours post invasion (hpi) and their day 6 gametocyte conversion rate (GCR) **(b)**. Dotted and continuous lines indicate Shld1 (-) or (+), respectively. **c**) Schizont stage *ap2-g* expression analysis from Shld1-regulated *gdv1* line with wt (*ap2-g.V//gdv1.fkbp*) or mutant (*ap2-g.L//gdv1.fkbp*) *ap2-g* and also wt (*ap2-g.V.fkbp// wt gdv1)* and mutant (*ap2-g.L.fkbp//wt gdv1*) *ap2-g* Shld1*-*regulated lines with wt *gdv1*. (+) or (-) indicate presence or absence of Shld1. **d-e)** *Ap2-g* and *msrp1* gene expression analysis in sorbitol synchronized ring stage parasites (6-10 hpi) in wt *ap2-g* parasite line (*NF54.gdv1.3xha.fkbp*) in the presence (GDV1(+)) or absence (GDV1(-)) of Shld1, *ap2-g* mutation corrected parasite line,*3D7.stk2Δ//ap2-g.V* (AP2-G.V), *ap2-g* mutated parasite lines, *NF54.ap2-g.L* (AP2-G.L), *3D7.stk2Δ//ap2-g.L* (B4_L_) and F12 with a premature stop codon in *ap2-g*. Each circle indicates an independent data point and the bar and error bar represent mean and standard error of the mean, respectively. A Kruskal-Wallis test followed by Dunn’s multiple comparison was used for statistical analysis between groups. The p value of each comparison is indicated by **** (<0.0001). **f**) Schematics showing *ap2-g* initial expression, autoregulation and AP2-G dependent downstream genes in the different engineered lines in the presence or absence of Shld1. Dotted circle indicates unknown potential co-factors, GTAC indicates the AP2-G binding DNA motif (GnGTAC), 1) depicts initial transcription, 2) depicts autoregulation that requires a functional AP2-G, and 3) depicts AP2-G dependent gene expression. Yellow circles indicate the FKBP domain that targets the attached protein for degradation in the absence of Shld1 ligand. Red or blue pentagons indicate AP2-G.V or AP2-G.L, respectively.

From the time course, 40-44 hpi was selected to compare *ap2-g* expression in lines with wt *gdv1* and wt or mutant *ap2-g* tagged with *fkbp* allowing Shld1-mediated regulation (*ap2-g.V.fkbp//wt gdv1*, *ap2-g.L.fkbp//wt gdv1*) as well as *fkbp*-tagged *gdv1* lines expressing non-tagged wt or mutant *ap2-g* (*gdv1.3xha.fkbp//ap2-g.V* or *gdv1.3xha.fkbp//ap2-g.L*). *Ap2-g* transcription at schizont stage was completely disrupted by GDV1 knockdown in the absence of Shld1 regardless of whether the line expressed wt or mutated *ap2-g* **(Fig 4c).** In contrast, in the *fkbp*-tagged *ap2-g* wt or mutant lines expressing wt GDV1, as well as a gametocyte-deficient line, F12, which expresses wt GDV1 but has a truncated AP2-G lacking the AP2 domain ^10^, *ap2-g* was still expressed (∼16 fold or more) in the presence or absence of Shld1. However, *ap2-g* expression was higher in *ap2-g.V.fkbp* line (Shld1+) than the *ap2-g.L.fkbp* line (Shld1+) or either line in the absence of Shld1. These results are consistent with wt GDV1 initiating *ap2-g* expression in all these lines, while only Shld1-stabilized wt AP2-G.V stimulated AP2-G mediated autoregulation. *Ap2-g* transcript levels were also higher in the presence of Shld1 in the *gdv1.fkbp*//*ap2.g.V* line than the *gdv1.fkbp*//*ap2-g.L* line (**Fig 4c**).

In the next generation, only parasites expressing both AP2-G.V and GDV1 [*3D7.stk2Δ//ap2-g.V*, *NF54-gdv1.3xha.fkbp//Ap2-g.V* (Shld1+)] expressed all of the AP2-G dependent early gametocyte genes ^4^ tested (**Fig 4d-e, Fig S4a-g**) that contain at least one copy of the upstream AP2-G DNA recognition motif, GnGTAC. In contrast, none of these early gametocyte genes were expressed by parasite lines expressing AP2-G.L (*3D7.stk2Δ//ap2-g.L*, *NF54.ap2-g.L*), truncated AP2-G (F12) or those lacking GDV1 [*NF54.gdv1.3xha.fkbp//ap2-g.V* (Shld1-)]. This clearly demonstrates that after GDV1 initiates *ap2-g* expression AP2-G.V, not AP2-G.L, autoregulates *ap2-g* during schizogony and stimulates downstream gene expression in the progeny consistent with the need for DNA binding to mediate these effects (**Fig 4f**).

### Quantification of *ap2-g.V & ap2-g.L* and *gdv1* translation

A nanoluciferase (nLuc) reporter was used to quantify and compare the time course of *ap2-g.V, ap2-g.L,* and *gdv1* translation using a series of transgenic lines (*3D7.stk2Δ//ap2-g.V.fkbp.p2a.nLuc, 3D7.stk2Δ//ap2-g.L.fkbp.p2a.nLuc* and *NF54.gdv1.p2a.nLuc*). The P2A peptide that causes ribosomal skipping ^27, 28^ was included upstream of nLuc to allow the production of two separate proteins from a single RNA. We first confirmed that the nLuc signal was linear through the parasitemias used in the experiments (**Fig S5**). The assay was then used to monitor nLuc activity from 20 to 44 hpi and again 9 & 22 hours later (53 hpi & 66 hpi) in the next generation rings. The time course data shows that signal from *gdv1*-linked nLuc was first detected at 26 hpi, which is prior to *ap2-g-*linked nLuc activity, as expected for an upstream regulator. *Gdv1*-linked nLuc activity peaked at 34 hpi and remained stable through schizont development (**Fig 5a**). This early GDV1 activity is also supported through live imaging of a reporter line (*683.gdv1.3xha.tdTom.fkbp*) that first detected GDV1-tdTomato fluorescent signal at 28-30 hpi which corresponds to the late trophozoite to early schizont transition prior to the first nuclear division **(Fig 5b)**. As schizogony progressed, GDV1 expression continued and a punctate pattern around each nucleus was observed (**Fig S6**). In contrast to *gdv1*-linked nLuc, *ap2-g.V-*linked nLuc activity was first detected at 31 hpi and continued to increase until 41 hpi after which activity plateaued, consistent with its transcription time-course (**Fig 4a**). The time course of *Ap2-g.L*-linked nLuc signal was similar to *ap2-g.V*-linked nLuc activity during schizogony but levels were 2-3 fold lower in magnitude than *ap2-g.V*-linked nLuc activity, even throughout the plateau at 39-44 hpi (**Fig 5a**). In the next generation ring stage parasites (53 & 66 hpi) *ap2-g.V*-linked nLuc activity remained high, which is in marked contrast with the *ap2-g.L-*linked nLuc line that had ∼12 fold less nLuc activity and no gametocyte conversion (**Fig 5a, fig S7**). The increased *ap2-g.V*-linked nLuc activity in comparison to *ap2-g.L*-linked nLuc activity in schizonts (38-40 hpi) and next generation rings stage parasites (10-15 hpi) was confirmed using additional *ap2-g.L* (*NF54.ap2-g.L.p2a.nLuc, 3D7.stk2Δ//ap2-g.L.fkbp.p2a.nLuc*) and *ap2-g.V* (*NF54.ap2-g.V.p2a.nLuc, 3D7.stk2Δ//ap2-g.V.fkbp.p2a.nLuc*) parasite lines with NF54 or 3D7 background. Consistently there was a significant increase in nLuc activity in schizonts (3-4 fold) and rings (4-10 fold) stages in *ap2-g.V* over *ap2-g.L* parasite lines with the same parent **(Fig 5c-d)** as well as a significant increase in gametocyte conversion rate in the *ap2-G.V* lines **(Fig. 5e)**. This pattern is consistent with an increase in AP2-G.V, not AP2-G.L, protein expression due to AP2-G.V-dependent autoregulation. The nLuc activity observed in *ap2-g.L* lines in schizont stage strongly supports wt GDV1 driving the initial *ap2-g.V* or *ap2-g.L* transcription, that is then augmented in the *ap2-g.V* lines due to autoregulation. The markedly lower levels of *ap2-g.L*-linked nLuc activity in the next generation rings is consistent with the decrease in GDV1-linked nLuc activity in rings (**Fig 5a**) as well as the lack of functional AP2-G.V to maintain expression in the absence of GDV1. The findings also highlight the importance of AP2-G.V expression in gametocyte maturation.

**Fig 5.**
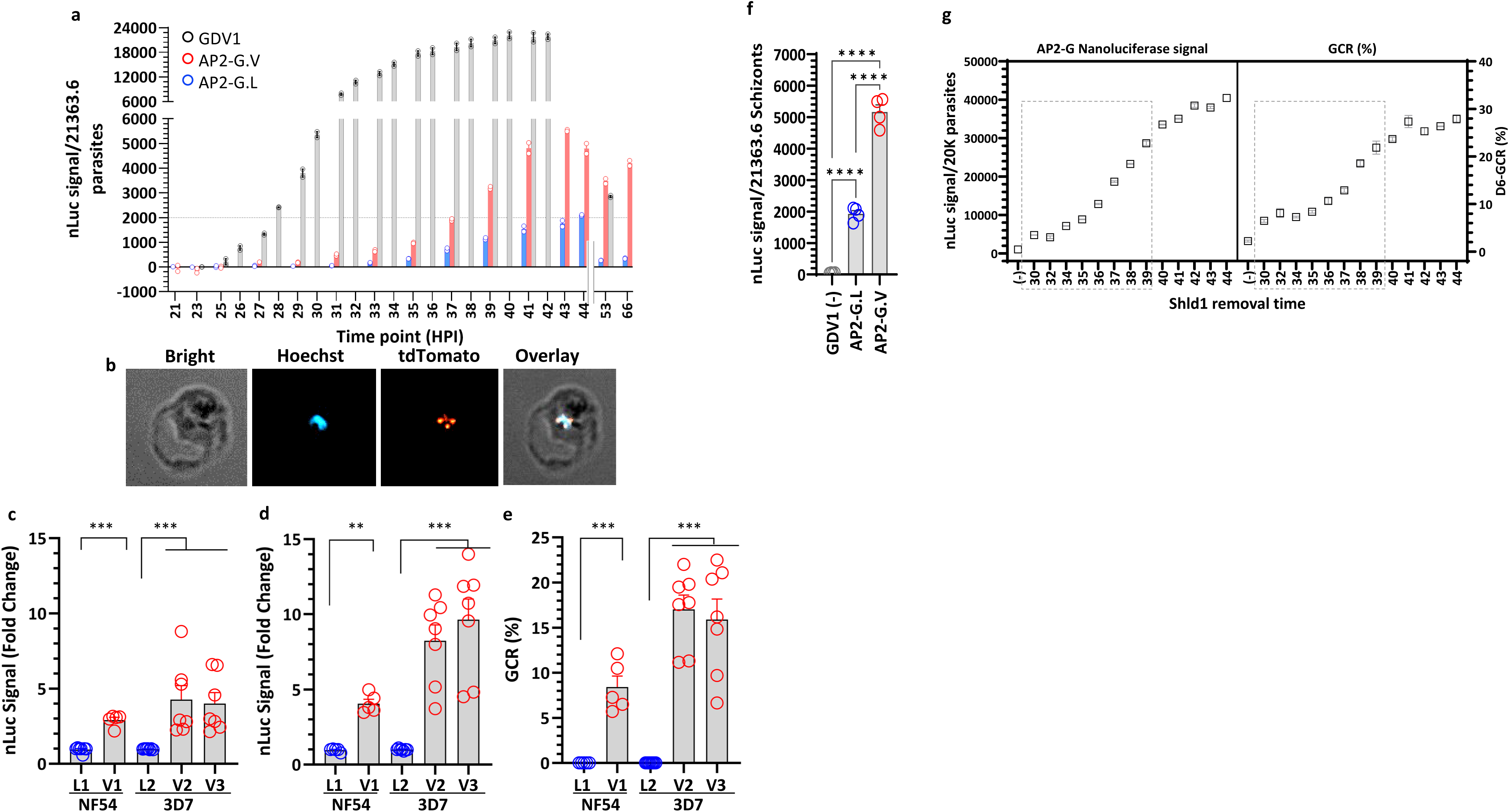
*Gdv1* and *ap2-g* reporter lines demonstrate that GDV1 is required for the initiation of *ap2-g* expression. **a)** Time course analysis of *ap2-g* and *gdv1*-linked nLuc activity from early trophozoites (20 hpi) to the next generation of rings (53, 66 hpi). nLuc activity was monitored in *ap2-g.V* (*3D7.stk2Δ//ap2-g.V.fkbp.p2a.nLuc), ap2-g.L* (*3D7.stk2Δ//ap2-g.L.fkbp.p2a.nLuc*) and *gdv1* (*NF54.gdv1.P2A.nLuc*) nLuc lines. **b)** Live imaging of GDV1.tdTomato expression using the *683.gdv1.tdTomato.fkbp* (Shld1+) line stained with Hoechst to localize the nucleus. Fold change in nLuc activity in 40-44 hpi schizonts **(c)** and in the next generation 5-10 hpi rings (**d)** and the corresponding gametocyte conversion rate (GCR) at D4 **(e)** in *ap2-g.L* [*NF5.ap2-g.L.p2a.nLuc* (L1) and *3D7.stk2Δ//ap2-g.L.fkbp.p2a.nLuc* (L2)] and in *ap2-g.V* [*NF54.ap2-g.V.p2a.nLuc* (V1) and *3D7.stk2Δ//ap2-g.V.fkbp.p2a.nLuc* (V2, V3)] parasite lines. **f**) *Ap2-g* linked nLuc signal from *NF54.gdv1.3xha.fkbp//ap2-g.V.p2Aa.nLuc* without Shld1 (GDV1-), *NF54.ap2-g.L.p2a.nLuc* (AP2-G.L)*, and NF54.ap2-g.V.p2a.nLuc* (AP2-G.V) reporter lines. **g)** GDV1 regulated *ap2-g-*linked nLuc activity and GCR quantification using the *NF54.gdv1.3xha.fkbp//ap2-g.V.p2a.nLuc* line. The X-axis indicates the time points when Shld1 was removed from the culture well and the dotted box indicates developmental phase that requires GDV1 for gametocyte production. Assay details are provided in **Fig S9b**. **c-f**: Red and blue circles indicate *ap2-g.V* and *ap2-g.L*, respectively. Each circle indicates an independent data point, and the bar and error bar represent mean and standard error of the mean, respectively. A Kruskal-Wallis test followed by Dunn’s multiple comparison was used for statistical analysis between groups. The p value of each comparison is indicated by ns (>0.05), * (<0.05), ** (<0.01), *** (<0.001), and **** (<0.0001).

To directly assess the timing of the requirement for GDV1 in *ap2-g* transcription initiation, we generated *NF54.gdv1.3xha.fkbp//ap2-g.V.p2a.nLuc* transgenic reporter that allows monitoring of *ap2-g* activity by regulating GDV1 with Shld1. As expected in the absence of Shld1 there was no *ap2-g.V*-linked nLuc activity (**Fig 5f**) and following the addition of Shld1 at ring stages (16hpi) *ap2-g.V*-linked nLuc activity began at ∼33 hpi and began to plateau at 40 hpi (**Fig S8a).** To determine when GDV1 was no longer needed for *ap2-g.V* expression, Shld1 was added to tightly synchronized cultures in a 24 well plate at 16 hpi and then removed from replicate wells at defined time points from 30 hpi to 44 hpi (**Fig S8b**). *Ap2-g.V-* linked nLuc activity was assessed in all wells at 44 hpi (schizonts) and 56 hpi (rings). The GCR of each culture was also measured **(Fig S8b).** Early GDV1 removal (<40 hpi) resulted in lower *ap2-g.V*-linked nLuc levels in both schizonts (44 hpi) and the next generation rings (56 hpi), as well as lower gametocyte production (**Fig 5g, Fig S8c**). Once parasites reached 40 hpi, *ap2-g.V*-linked nLuc levels were maximal and removing GDV1 had little effect (**Fig 5g**). This controlled GDV1 exposure indicates it is required until 39 hpi, for maximal *ap2-g.V* expression during the trophozoite-schizont transition.

### Dynamics of AP2-G autoregulation and downstream gene induction

*Ap2-g.mScarlet* (mSc) reporter lines (*3D7.stk2Δ//ap2-g.L.mSc* or *3D7.stk2Δ//ap2-g.V.mSc*) were generated to compare AP2-G.V and AP2-G.L protein expression patterns and identify sexually committed schizonts. Live imaging and flow cytometry show a subpopulation of schizonts (9-11%) positive for AP2-G.V.mSc which is similar to the 8-10% day 4 GCR (**Fig 6a**, **Fig S9a-b**), suggesting the positive schizonts progressed to gametocytes. In contrast, none of the *ap2-g.L.mSc* lines were positive and there was no GCR (**Fig 6a, Fig S9a-b**). It is possible that the fluorescence assays are not sensitive enough to detect the amount of AP2-G.L generated from the low transcript levels induced by GDV1 in the absence of AP2-G autoregulation or, alternatively, that the DNA binding activity of AP2-G.V is required for protein expression.

**Fig 6.**
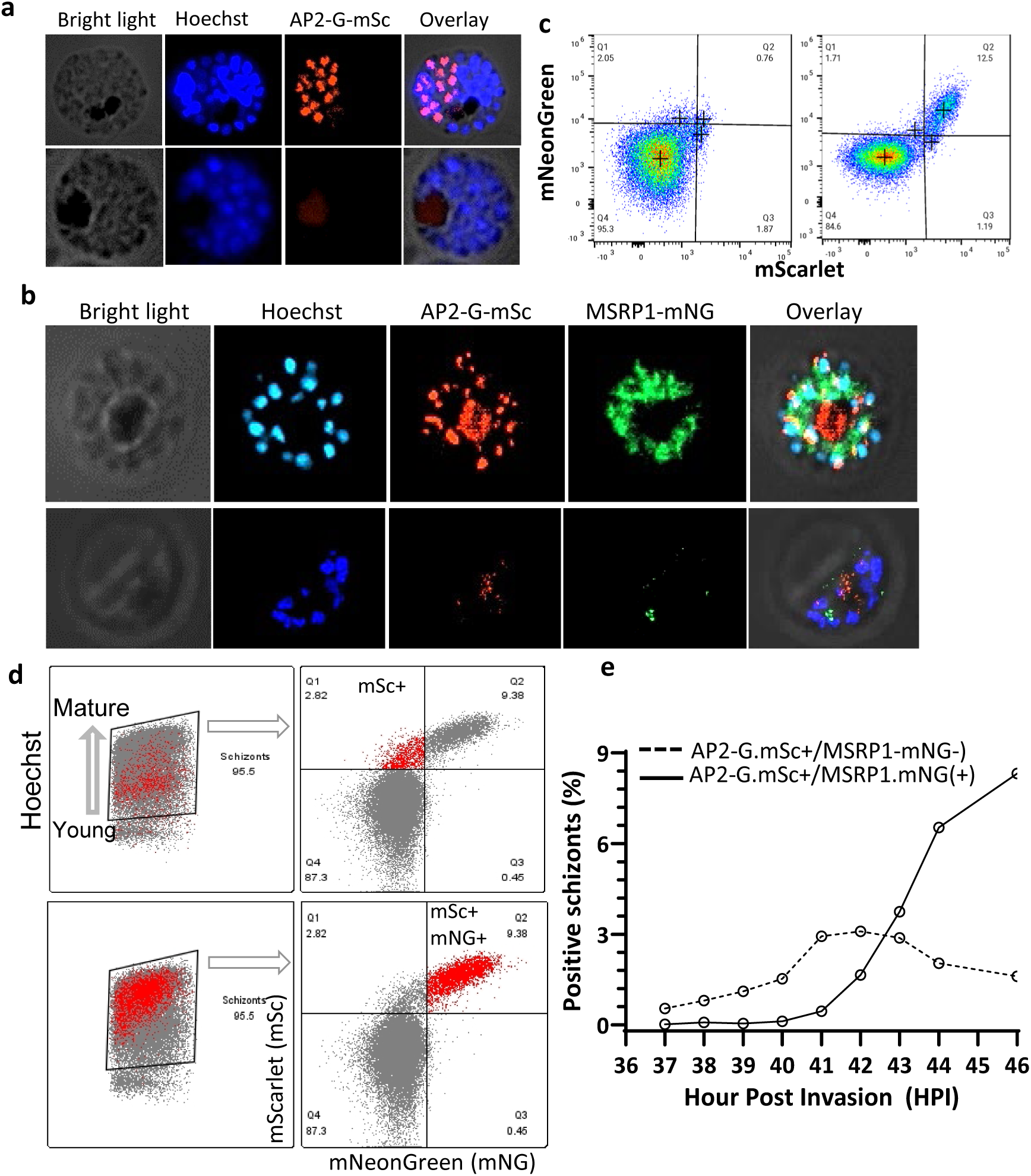
Live imaging demonstrates the timing of AP2-G.V dependent autoregulation and MSRP1 gene expression. **a)** Live imaging showing expression of AP2-G.mScarlet (mSc) in a schizont stage parasite from mutated (*3D7.stk2Δ//ap2-g.L.mSc*: bottom panel) or mutation corrected (*3D7.stk2Δ//ap2-g.V.mSc*: top panel) lines that were stained with Hoechst to detect DNA. **b)** Live imaging of double reporter lines showing AP2-G and MSRP1 expression in the mutation corrected line (*3D7.stk2Δ//ap2-g.V.mSc//msrp1. mNeonGreen (mNG)*: top panel), not the mutated line (*3D7.stk2Δ//ap2-g.L.mSc//msrp1.mNG*: bottom panel). Both parasite lines were stained with Hoechst to detect DNA. **c**) Quantification of AP2-G.mSc and MSRP1.mNG positive schizonts in the mutated (**left panel**) and mutation corrected (**right panel**) lines shown in **b**. **d**) Retro analysis of flow cytometer data shows AP2.G.mSc positive schizonts (top panel) have less Hoechst staining, consistent with younger schizonts with less DNA, whereas schizonts positive for both AP2.G.mSc and MSRP1.mNG (bottom panel) have higher Hoechst staining suggesting they are older. **e**) Flow cytometric quantification of single positive (AP2-G.mSc+/MSRP1mNG-: dotted line) or double positive (AP2-G.mSc+/MSRP1.mNG+: continuous line) schizonts from 37 to 46 hpi using *3D7.stk2Δ//ap2-g.V.mSc//msrp1.mNG* parasites. Details of flow cytometric quantification are provided as **Fig S11**.

To extend the expression profile of sexually-committed schizonts beyond *ap2-g*, the RNA time course of *msrp1* (ID# PF3D7_1335000) another gene expressed in sexually committed schizonts was tested. *Msrp1* transcript was only detected in AP2-G.V line and the time course was slightly later than *ap2-g* RNA, (**Fig S10a**) consistent with earlier reports that it is AP2-G regulated ^3, 4, 12^. Protein expression was tracked by tagging MSRP1 with mNeon Green (mNG) in 3D7 and NF54 parental lines (*3D7.stk2Δ//ap2-g.L.mSc//msrp1.mNG*, *3D7.stk2Δ//ap2-g.V.mSc//msrp1.mNG*, *NF54.msrp1.mNG//ap2-g.V.mSc* and *NF54.msrp1.mNG//ap2-g.L.mSc*). AP2-G.V.mSc and MSRP1.mNG were co-expressed in a subpopulation of schizonts, while signal from neither protein was detected in the AP2-G.L lines using fluorescence microscopy or flow-cytometry. AP2-G.V.mSc expression was punctate and co-localized with the nuclear stain, while in mature schizonts MSRP1 was concentrated around the periphery of the merozoites (**Fig 6b (top panel), Fig S10b**). AP2-G^+^MSRP1^+^ double positive schizonts were found only in the *AP2-g.V* lines and they were in the same proportion as the D4 GCR (**Fig 6b-c, Fig S10d-e**) and a similar correlation was found between MSRP1^+^ schizonts and D4 GCR in the *NF54.msrp1.mNG* line (**Fig S10b-c**) suggesting MSRP1 can be used as another marker of sexual commitment.

We also found that MACS-purified schizonts from a dual reporter line (*NF54.ap2-g.V.mSc//msrp1.mNG*) that were only positive for AP2-G.mSc, not MSRP1.mNG, had less DNA, suggesting they were younger, than schizonts double positive for AP2-G.mSc and MSRP1.mNG (**Fig 6d**). In these dual reporter lines there were also no MSRP1.mNG single positive schizonts, only double positive AP2-G.mSc / MSRP1.mNG schizonts, consistent with MSRP1 expression being dependent on AP2-G. The time course of AP2-G.V.mSc and MSRP1.mNG (*NF54.ap2-g.V.mSc//msrp1.mNG)* expression was further evaluated every hour from 28-46 hpi to understand the dynamics of sexual commitment. AP2-G.mSc single positive schizonts were first observed from 39-40 hpi, typically in 4-5 nuclear schizonts, and none of the schizonts with less than three nuclei were positive. Approximately 2 hrs later (41-42 hpi), these AP2-G positive schizonts turned positive for MSRP1.mNG **(Fig 6e, Fig S11**). Again, no single positive MSRP1.mNG schizonts were detected. The detection of AP2-G and MSRP1 expression was much later than GDV1.tdTom, which was detected prior to nuclear division at 28-29hpi (**Fig 5b, Fig S6**). These distinct MSRP1.mNG expression patterns allow differentiation of early and late sexually committed schizonts and demonstrates the distinct dynamics of AP2-G.V and downstream gene induction.

## Discussion

A single valine to leucine mutation at the beginning of the AP2 DNA binding domain in AP2-G completely blocked DNA binding and gametocyte production demonstrating the critical role of this AP2-G transcription factor for sexual differentiation. In contrast, we found the initiation of *ap2-g* expression during early schizogony requires GDV1, not functional AP2-G. Whereas, later in mature schizonts and the next generation ring stages AP2-G.V-dependent auto-upregulation plays a major role in maintaining *ap2-g* expression and driving downstream gene expression. These regulatory differences allowed the development of stage specific reporters that differentiate the initial induction of *ap2-g*, early AP2-G autoregulation followed in 2 hours by the expression of the downstream gene, *msrp1.* The lines provide tools to further assess the underlying molecular mechanisms and screen for factors regulating malaria transmission.

Our detailed assessment of the time-course of sexual commitment demonstrates that *gdv1* translation is first detected at 26 hpi and plateaus at 34 hpi, prior to the initiation of both wt and mutant (L) *ap2-g* transcription and translation at 30-31 hpi. The early expression of GDV1 is consistent with its proposed role in de-repressing the *ap2-g* locus that has been epigenetically silenced by H3-K9Me3 and HP1 binding upstream of the *ap2-g* promoter ^11, 14^. The role of GDV1 prior to AP2-G autoregulation is also supported by the almost complete block in *ap2-g (V* and L) transcripts when GDV1 is knocked down in the absence of Shld1. In contrast, *ap2-g* transcripts are still generated in GDV1 expressing lines in the absence of functional AP2-G, either due to knock down or changing AP2-G V to L. Defining this early period of *ap2-g* activation prior to the requirement for AP2-G.V expression can be used to focus further investigation of the mechanisms involved in de-repression, such as HP1 and histone 3 lysine 9 trimethylation removal and the participation of additional players to fully understand the initiation of sexual commitment in a subpopulation of parasites. AP2-I ^12, 29^, AP2-G5 ^30^ and a recently identified AP2-P ^31^ have been reported to bind upstream of *ap2-g* and could also contribute to its regulation.

After 38 hpi, the fluorescent detection of AP2-G.V coupled with the higher expression level of *ap2-g* in AP2-G.V expressing lines in comparison to lines lacking AP2-G, or those expressing AP2-G.L or truncated AP2-G suggests the expression increase requires functional AP2-G and is likely due to autoregulation. Additionally, during this phase (>38 hrs), knocking down GDV1 had little effect on *ap2-g.V-*linked nLuc expression or sexual conversion. AP2-G.V is also required for MSRP1.mNG expression in sexually committed schizonts and allows separation of AP2-G.V.mSc single positive parasites prior to MSRP1.mNG expression to evaluate early changes associated with AP2-G.V autoregulation. The effect of AP2-G.V autoregulation in maintaining *ap2-g* transcription is even more pronounced in the next cycle rings and early stage I gametocytes and suggests AP2-G.V is likely to be the primary activator of continued sexual differentiation and downstream gene transcription. In *P. berghei*, AP2-G is primarily detected in a subpopulation ring stage, not schizonts, that progressed to gametocyte production, not asexual replication, consistent with the importance of AP2-G in early gametocyte formation ^32, 33^. In addition to demonstrating the importance of the V_2163_ to AP2-G function and dissecting the roles of GDV1 and AP2-G.V in *P. falciparum* sexual commitment, these lines provide sensitive tools to directly quantify sexual commitment in schizont stages and allow small molecule library screens ^34, 35^ for compounds that block sexual differentiation. Additionally, the inability of AP2-G.L to stimulate AP2-G dependent genes expression, allows the mutated lines to be used to focus the analysis on the regulation of the *ap2-g* locus without interference from AP2-G autoregulation or downstream gene expression. Both fluorescent and nLuc reporter lines can also be used to screen plasma/serum samples to identify human factors that regulate sexual conversion which vary dramatically in malaria patients ^3, 4, 25^.

In contrast to the essential role of *gdv1* and *ap2-g* in sexual differentiation*, pfstk2* appears to be dispensable for asexual or sexual growth. This finding is consistent with previous piggyBac transposon-based insertion mutagenesis ^23^ in *P. falciparum* monitoring asexual growth and the analysis of sexual development following knockout *P. berghei* ^36^. Interestingly, *pbstk2* which is expressed in mosquito and liver stages, has been shown to play a critical role in the maturation of liver stage schizonts ^36^. The *pfstk2*-*KO* or inducible transgenic lines expressing AP2-G.V generated in the course of our work could be used to investigate the role of *pfstk2* in *P. falciparum* liver stage development.

The functional importance of a single valine located at the beginning of the predicted AP2 DNA recognition domain of AP2-G is clearly demonstrated by this work. Simply changing valine to leucine, a similar hydrophobic amino acid, blocked gametocyte production, the auto-upregulation of *ap2-g* transcription, translation and protein production as well as the up-regulation of downstream genes expressed in sexually committed schizonts or the next generation of ring stage parasites. The critical nature of this site in AP2-G is further supported by in silico analysis predicting an alteration in the AP2 DNA recognition site when V_2163_ is replaced by any other amino acid and more directly by the inability of recombinantly expressed AP2 domain of AP2-G. V_2163_L (aa 2150-2220) to bind an oligonucleotide containing the AP2-G binding motif, GnGTAC. We also found that the same V_2163_L mutation was present in some previously reported *P. falciparum* gametocyte non-producer piggyBac mutants, including clone IGM 2E4 ^37^. That study also used 3D7 as the parental strain, making it likely that AP2-G-V_2163_L is a pre-existing mutation already present within a sub-population of 3D7. Together the data strongly suggests changing valine to leucine at position 2163 disrupts DNA binding thereby blocking the ability of AP2-G.L to activate transcription of itself or downstream genes and abrogating gametocytogenesis.

The lack of missense mutations among the thousands of clinical isolates sequenced to date (PlasmoDB^22^, MalariaGEN^26^) also supports the importance of V_2163_ in sexual differentiation, which is required for person-to-person transmission through the community. Conversely, in vitro culture depends only on asexual growth for propagation. Therefore, in vitro a mutation that inhibits ∼10% of schizonts from committing to gametocytes would increase asexual growth by ∼10% providing selective pressure for the mutant parasite in in vitro culture. This in vitro growth advantage of sterile parasite lines always has to be considered when screening transgenic lines and highlights the importance of complementation and whole genome sequencing when analyzing phenotypes. There are multiple examples which suggest that gametocytogenesis phenotypes were selected for in vitro cultures, including deletion of parts of chromosome 9 containing *gdv1* ^9^ or mutations causing premature termination of GDV1 ^24, 25^ or AP2-G ^10^. Interestingly, as opposed to all other known mutation/deletions conferring gametocyte deficiency through *gdv1* or *ap2-g*, V_2163_L is the first mutation that alters AP2-G function without causing premature termination consistent with the key role played by V_2163_ in protein function. Outside the AP2 domain of *ap2-g* there are a large number of mutations in *ap2-g* in worldwide geographical isolates (PlasmoDB ^22^, MalariaGEN^26^) that have not been functionally evaluated.

In *Plasmodium* a conservative valine to leucine substitution with a strong phenotype, such as drug resistance, is rare but not unprecedented. A V to L mutation has been reported for *P. falciparum cytochrome b* where V259L is selected with atovaquone drug pressure in multiple studies ^38, 39, 40, 41^ and in *P. vivax dhfr* V64L is associated with antifolate resistance in clinical isolates ^42^. A loss of gene function due to a single valine to leucine mutation has been also demonstrated in various organisms such as a V875L in *cifb* reduces cytoplasmic incompatibility in Wolbachia bacteria ^43^, a V865L substitution in a human androgen receptor reduces sensitivity to androgen hormone ^44^, and a V321L substitution in *neuregulin1* in the hippocampus disrupts its nuclear localization resulting in a complete loss of the function ^45, 46^.

In summary, the data strongly supports the vital role of valine_2163_ in the AP2 domain of AP2-G for DNA binding, sexual commitment and early gametocytogenesis. In contrast, GDV1 is required for the initial activation of *ap2-g* transcription from 30-39 hpi and functional AP2-G during this phase is not essential. The definition of the time course and the development of lines with inducible GDV1 expression in the presence and absence of functional AP2-G allows the direct evaluation of the signaling pathways involved in *ap2-g* activation. AP2-G and MSRP1 reporter lines also clearly differentiate between early and late stages of sexual commitment during schizogony providing new tools to assess the effect of factors to modulate gametocyte conversion and transmission through the community.

## Materials and Methods

### Parasite culture

*Plasmodium falciparum* parasite strains (NF54, 3D7) or transgenic or mutant lines, were cultured in a complete RPMI medium containing RPMI 1640, 25 mM HEPES, 100 μg/ml of hypoxanthine, and 0.3 mg/ml of glutamine (KD Biomedical, Columbia, MD) supplemented with 25 mM NaHCO3 (pH 7.3), 5 μg/ml of gentamicin, and 10% human serum (Interstate Blood Bank, Memphis, TN) or 0.5% Albumax II (Gibco, USA) in an atmosphere of N_2_:O_2_:CO_2_ 90:5:5. Sorbitol treatment (5%, 10–30 min at 37 °C) or MACS® LS columns (Miltenyi Biotec, Auburn, CA) were used to synchronize parasites in ring or schizont stages, respectively. Parasitemia [(parasite-infected RBCs/total RBCs) *100] was evaluated through microscopy of culture smears stained with 10% Giemsa for 15 m (Sigma-Aldrich, St Louis, MO).

### *P. falciparum* gametocyte phenotyping

The gametocyte conversion rate (GCR) from a single cycle was determined by quantifying parasitemia of a synchronized ring stage culture (Day 0) then adding N-acetylglucosamine (50 mM) (NAG) to block further asexual growth. The plate/flask was then cultured for five or seven days (D4 or D6) with daily media changes without RBC supplementation and the D4 or D6 gametocytemia (stage II-III or III-IV) determined **(Fig S1d)**. Parasites were quantified using Giemsa stained culture smears or flow cytometry, which is described in the next section. The GCR of each sample was calculated using the formula (D4 or D6 gametocytemia / D0 ring parasitemia) x 100.

### Flow cytometry and data analysis

Parasite cultures and uninfected RBCs (negative control) were resuspended at 0.02% hematocrit (∼2,000 cell/µl) in 1x SA buffer (154 mM NaCl, 9.27 mM glucose, 10 mM Tris-HCl, pH 7.4) with 500 nM MitoProbe DilC_1_(5) (Cat: M34151) and incubated at 37°C in the dark for 30 min prior to analysis on an Attune Nxt flow cytometer (Invitrogen, USA). The forward and side scatter signals were used to select the total RBC population and then the DilC_1_(5) (Ex 640 nm/Em 675 ± 25 nm) was used to identify *P. falciparum* infected RBCs or Pf infected RBC.

Cytek Aurora Spectral Analyzer flow cytometer (USUHS, Biomedical Instrumentation Center facility) was used to identify parasite infected RBC positive for mScarlet (mSc) or mNeonGreen (mNG) or both fluorescent protein signals from transgenic lines. Hoechst 33342 DNA stain (Invitrogen, USA, Cat: H3570), 1:500 dilution, 10 minutes staining) was used to quantitate and identify the stage of infected RBCs (iRBCs). The proportion of mSc and or mNG positive iRBCs observed in total or stage-specific iRBCs was determined using FlowJo v10.10.0. A retro gating strategy was used to visualize and quantify only AP2-G.mSc positive iRBCs or MSRP1.mNG positive iRBCs from MACS-column purified parasites having a range of younger to mature schizonts based on the Hoechst 33342 signal.

### Plasmid engineering (CRISPR & non-CRISPR) and transgenic line generation

The *stk2* knockout lines (*3D7.stk2Δ.wr* or *NF54stk2Δ.wr*) were generated using a double crossover plasmid (*pCC1*) in the absence of Cas9. To generate this plasmid, two homologous fragments were introduced in the plasmid using *Sac*II + *Spe*I and *Nco*I+*Avr*II restriction enzymes. T4-DNA ligase was used to ligate the restriction enzyme-digested PCR products into the plasmid. The inducible *stk*2 line was generated using a single crossover plasmid (*p1605.stk2.gfp.fkbp.wr*) (**Fig S12a**) to tag the C-terminus of *stk2* with green fluorescent protein (GFP) followed by the FKBP-destabilization domain (FKBP) ^3^. This single crossover plasmid was also used to generate an *ap2-g* mutated line (*NF54.ap2-g.L.gfp.fkbp.wr*). To generate the single crossover plasmid construct, a single homologous region (600-1500 bp) of the target gene was PCR amplified, digested with XhoI + AvrII restriction enzyme and ligated into XhoI + AvrII digested plasmid using T4-DNA ligase. Primer DNA sequences for *stk2* and *ap2-g* genes are given in supplementary **table S2**.

*E. coli* electrocompetent cells (DH10B) were transformed with the ligation reaction and grown overnight on ampicillin (100 µg/ml) containing LB plates. Ampicillin-resistant insert positive colonies were assessed using restriction enzyme digestion or by PCR and then further verified by DNA sequencing. DNA sequence-confirmed colonies were expanded and plasmid was purified using the NucleoBond Xtra Plasmid Maxi prep kit (Macherey Nagel, USA). Purified plasmid reconstituted in 1x cytomix buffer containing KCl (120mM), CaCl2 (0.15mM), K2HPO4/KH2PO4 (10mM, pH 7.6), HEPES (25mM, pH 7.6), EGTA (2mM, pH 7.6), and MgCl2 (5mM) and stored in −20°C. 100–150 µg plasmid DNA per transformation was used to transform parasites using established transformation protocols ^3^. Approximately 5% rings were transformed (Day 0) and on D2 appropriate drug was added for selection, WR99210 (2.5nM) (Jacobus Pharmaceuticals, Plainsboro, NJ) or Blasticidin (2.5 nM) (Gibco, USA). Resistant parasites from three independent transformations were selected and grown for 3 weeks without drug before reapplying the appropriate drug. The resistant parasites obtained were screened for the presence of a single or double-crossover chromosomal integration **(Fig S1b, S12b)** using PCR with primers listed in **table S2** and selected for cloning by limiting dilution. Transformed lines were maintained in appropriate drugs.

CRISPR/Cas9 gene editing was used to generate the majority of the *gdv1, ap2-g* and *msrp1* transgenic lines. A single plasmid (*pDC-cam-cas9-gfp.wr*) ^47^ or two-plasmid based CRISPR/Cas9 gene editing was employed to transform gametocyte competent laboratory strains (3D7, NF54) or gametocyte deficient precursor transgenic lines (*3D7.stk2Δ-WR*, *NF54.ap2-g-L-gfp-fkbp-WR*). *pDC-cam-cas9.gfp.wr* has a repair template (Flanked by EcoRI+AatII), a guide RNA sequence and Cas9 expression cassettes ^47^ and this plasmid is also called as single-guide plasmid (SGP). We used this SGP plasmid to replace *ap2-g.V_2163_* with *ap2-g.L_2163_* (*pDC-cam.ap2-g.L_2163_.wr*) or vice versa *ap2-g.L_2163_* with *ap2-g.V_2163_* (*pDC-cam.ap2-g.V_2163_.bsd*), and also to tag the C-terminus of *gdv1* with *3xha.fkbp* (*pDC-cam.cas9.gdv1.3xha.fkbp-WR*) or *p2a.NanoLuc* (*pDC-cam.cas9.gdv1.p2a.NanoLuc-BSD)* or *tdTomato.fkbp* (*pDC-cam.cas9.gdv1.tdTomato.fkbp -BSD*), and the *msrp1* locus with *mNeonGreen* (*pDC-cam.cas9.msrp1.mNeonGreen-DSM*) **(Fig S12c-d)**. To tag c-terminus of *ap2-g* with a number of different reporters, we modified the SGP and generated a two-guide plasmid (TGP) by replacing *Cas9* with a 2^nd^ gSeq cassette using infusion cloning **(Fig S12d-e)**. Using this TGP, we tagged the c-terminus of *ap2-g* locus with *fkbp* (*pDC-cam.ap2-g(L or V).fkbp*), *mScarlet* (*pDC-cam.ap2-g(L or V).mSc*), *fkbp.p2a.NanoLuc* (*pDC-cam.ap2-g (L or V).fkbp.p2a.NanoLuc*) or *p2a.NanoLuc* (*pDC-cam.ap2-g(L or V).p2a.NanoLuc*). The WR drug resistance marker was replaced with either BSD or DSM1 depending on the requirement. When TGP was used for the transformation, if the parasite line used for transformation lacked *Cas9* expression a second plasmid *pUF-DSM* ^48^ was used as the *Cas9* source.

Guide RNA sequence (20 nt) with NGG PAM were designed using e-CRISP web-based tools ^49^ that provide only a few but very efficient guide sequences per gene. The gRNA seq was digested with *Bbs*I, a type IIS restriction enzyme, ligated into the plasmid using T4-DNA ligase and Stellar competent cells were used for the transfection. All other modifications (single or multiple donor fragments, drug resistance markers or the second gRNA cassette) were done using infusion cloning. To use this system, PCR fragments generated with infusion cloning compatible primers and then infused with the appropriate restriction enzymes digested plasmid. Insert positive colonies were identified and the NucleoBond Xtra Plasmid Maxi kit was used to purify plasmid for transformation as described above in the non-CRISPR plasmid section. The DNA sequences of the InFusion PCR primers and gRNAs used are listed in **table S2**. Fresh, uninfected RBCs were transformed using 100-150 ug of each plasmid, washed with RPMI, mixed with 0.2-1% MACS purified schizont stage parasites and incubated in triplicates in a 12 well culture plate. On D3 (ring stage) appropriate drug/s were applied for three generations/cycles (∼ 6 days) to select transformed parasites and then removed. Parasites recovered after drug/s selection were tested for editing using PCR (**Fig S12c**) and confirmed by DNA sequencing. Transformed parasites were cloned after genotype verification using a limiting dilution method and clones were further tested by DNA sequencing.

### Quantitative reverse transcriptase-polymerase chain reaction (RT-qPCR)

RNA was extracted from parasites preserved in 1x RNA Shld1 (Zymo Research, USA) using RNA Micro kit (Zymo Research, USA), as per the manufacturer’s instructions. RNA was eluted in 36 µl RNAse-free water and purified RNA (4.0 µl or 8.0 µl) was additionally treated with ezDNAse™ before cDNA synthesis using SuperScript™ IV VILO™ Master Mix (Invitrogen, USA). A 1:20 dilution of cDNA was made and 2 µl from this dilution was used for each RT-qPCR well. RT-qPCR (QuantStudio7 Pro) was performed using gene-specific primers designed using the web-based tool Primer3Plus ^50^ (**Table S2**) and Fast SYBR Green PCR Master Mix (Applied Biosystems, USA) as described earlier ^3^. Primer efficiency was tested by serial dilution and ranged from 85 to 105%. All samples were run in duplicate and tested for both the gene of interest and the control constitutive gene, *arginyl tRNA* (schizont stage) or skeleton binding protein 1 (*sbp1*) (ring stage) on the same plate. Data was analyzed using QuantStudio™ Design & Analysis Software v2.6.0, and the ΔCt values were determined by subtracting the mean cycle threshold (Ct) value for *arginyl tRNA* or *sbp1* from the mean Ct of each test gene. The relative quantity (2^-ΔΔCt^) was calculated for each gene using RNA from *NF54-gdv1-3xha-fkbp* parasites grown in the absence of Shld1 or gametocyte deficient lines (*3D7.stk2Δ or NF54-ap2-g-L*).

### Whole genome sequencing

Whole Genome Sequencing was conducted using a short-read workflow. Genomic DNA was used as input for library preparation after mechanical shearing and subsequently end-repair, a-tailing and ligation with unique dual index adapters. Sequencing libraries were assessed for size distribution and absence of adapter dimers before sequencing on an Illumina NovaSeq 6000 using a 151+8+8+151 bp read parameter for run conditions. An average 26,329,252 sequences per samples were generated and subsequently mapped using the Burrows-Wheeler Alignment ^51^ against *P. falciparum* 3D7 reference genome (PlasmoDB)^22^. The average percentage of mapped reads per isolate was 78% and an average 74x coverage was achieved across the genome. The Genome Analysis Toolkit ^52^ was then used to identify single nucleotide polymorphism (SNP) variants per isolates (**Table S1**).

### In-silico prediction of AP2 domain and structural visualization of AP2-G-DNA interaction

MOTIF search ^53^, a web-based tool was used to predict the *in-silico* AP2 domain. This web tool has definitions of known protein domains, including AP2 domains derived from plants to higher eukaryotes. We provided wild type or mutated amino acid sequences of AP2-G as an input and for each sequence an AP2 domain was predicted with a significance score.

### Electrophoretic mobility shift assay

*Plasmodium falciparum* (Pf-AP2-G-L_2163,_ PfAP2-G.V_2163wt_) and *P. berghei* (PbAP2-G_wt_) recombinant proteins were prepared as described earlier ^54^. Electrophoretic mobility shift assay (EMSA) was performed using Light Shift EMSA kits (Thermo Scientific) ^10^ using 2 μg of protein and 20 fmol of biotinylated probe, as previously described ^54^. Briefly, probes and recombinant DNA-binding domains were incubated at RT for 20min before loading on a 0.5x TBE-buffered 6% polyacrylamide gel. Complexes were separated at 100V until the dye front had migrated 3/4 of the gel length. Probes were then transferred overnight onto Nylon membranes, immobilized by UV (302 nm) cross-linking for 15min, and visualized using strepdavidin-HRP conjugate and luminol detection solution (BioRad).

### Parasite live imaging

MACS-column purified schizont stage parasites were used for visualizing mScarlet or mNeonGreen or tdTomato fluorescent protein. Live parasites re-suspended in 1xSA were stained with Hoechst 33342 DNA stain (1:500 dilution, 10 m) and washed once in 1xSA before imaging. Live imaging was performed using Zeiss inverted fluorescence microscope (Axio Observer 7) under 630x magnification and ZEN v3.0 (blue edition) in-built analysis software.

### Protein quantification using nLuc system

Nanoluciferase (nLuc) bioluminescence signal was measured using a Nano-Glo® Luciferase Assay System (Cat# N1110, Promega, USA) using the Cytation 5 Biotek imaging system as per manufacturer’s instruction. Tightly synchronized parasite cultures (Sorbitol-MACS-sorbitol) were used to measure nLuc signals at defined time points or developmental stages (schizonts or rings). Nano-substrate was used in 1:50 dilution in the assay buffer (30 µl) with an equal volume of culture (30 µl) making final dilution of 1:100. Bioluminescence signals were normalized with equal number of parasites in each comparison group and number of parasites were counted using a flow cytometry or hemocytometer. In the time-course experiment, a background signal subtraction was applied. Mean nLuc signal obtained from 16-22 hpi was subtracted from each data-point measured later from 24-44 hpi. In generation 2, rings stage nLuc signal was compared between samples containing the same number of parasites without a background subtraction. For the fold change analysis (**Fig 5c-d**), nLuc signal was presented relative to nLuc signal observed in *ap2-L_2163_* line.

### Data analysis

GraphPad Prism 10.00 statistical package ^55^ was used to evaluate differences between parasite lines. All comparisons between wt and *stk2Δ* clones, Shld1 (+) and Shld1 1 (-) or *ap2-g-L_2163_* and *ap2-g-V_2163_* groups were analyzed using a Kruskal-Wallis test followed by Dunn’s multiple comparison. The p value of each comparison is indicated by ns (>0.05), * (<0.05), ** (<0.01), *** (<0.001), and **** (<0.0001). Correlations between groups or conditions were assessed using Microsoft excel to determine the Pearson correlation coefficient.

### Plasmids and parasite lines source

*P. falciparum* wt (Nf54, 3D7) and mutant F12 strains were obtained from B.E.I Resources (USA) and Prof Manuel Llinas’s lab (Pennsylvania State University, USA), respectively. Plasmid with *tdTomato* (pGEMT-PT2A-iRFP670-Tdtomato-GFP: Addgene plasmid # 111817) ^28^ or *mScarlet* (pmScarlet_C1: Addgene plasmid # 85042) ^56^ *mNeonGreen* (ER-mNeonGreen: Addgene plasmid # 137804) ^57^ or *NanoLuciferease* (pUAS-NanoLuc: Addgene plasmid # 87696) ^58^ were obtained from Addgene Inc (USA).

## Acknowledgements

This investigation received financial support from Public Health Service grants from the National Institute of Allergy and Infectious Diseases (NIAID), National Institutes of Health (NIH), KCW (R01 AI069314 and R01 AI103638) and ML [R01 AI076276 and R01-AI125565 with support from the Center for Quantitative Biology (P50 GM071508)], the Intramural Research Program of the NIAID, NIH (BJM) and intramural research funds from the Uniformed Services University of the Health Sciences, USA (KCW: C073384316). We thank Drs. Kateryna Lund and Sintayehu Gebreyohannes (USU-BIC facility) for flow cytometry support. We extend thanks to former lab members Christen Dona and Van Anh Nguyen for their support generating a transgenic line (*683.gdv1.3xha.tdTom.fkbp*) as well as Will Valiant for preparing a plasmid (*p1605.stk2.gfp.fkbp.wr*). The single plasmid CRISPR system (*pDC-cam-cas9-gfp.wr*) was a gift from Prof Marcus Lee (University of Dundee, UK). All Addgene plasmids containing fluorescent protein were gifts from the Li Qian lab (UNC Chapel Hill, USA, tdTomato), Dorus Gadella Lab (University of Amsterdam, Netherlands, mScarlet and mNeonGreen) and Robert Campbell Lab (University of Tokyo, Japan, NanoLuciferase). The authors declare no competing interests for themselves or their family members. The opinions and assertations expressed herein are those of the author(s) and do not reflect the official policy or position of the Uniformed Services University of the Health Sciences (USUHS), or Henry M. Jackson Foundation for the Advancement of Military Medicine, Inc. (HJF), or the Department of Defense (DOD).

## Notes

### Competing Interest Statement

The authors have declared no competing interest.

## References

1. WHO. WHO Reports.) (2024).

2. Stewart LB, Freville A, Voss TS, Baker DA, Awandare GA, Conway DJ. Plasmodium falciparum Sexual Commitment Rate Variation among Clinical Isolates and Diverse Laboratory-Adapted Lines. Microbiol Spectr 10, e0223422 (2022).

3. Usui M, et al. Plasmodium falciparum sexual differentiation in malaria patients is associated with host factors and GDV1-dependent genes. Nat Commun 10, 2140 (2019).

4. Prajapati SK, et al. The transcriptome of circulating sexually committed Plasmodium falciparum ring stage parasites forecasts malaria transmission potential. Nat Commun 11, 6159 (2020).

5. Hofmann N, Mwingira F, Shekalaghe S, Robinson LJ, Mueller I, Felger I. Ultra-sensitive detection of Plasmodium falciparum by amplification of multi-copy subtelomeric targets. PLoS Med 12, e1001788 (2015).

6. Geiger C, et al. Declining malaria parasite prevalence and trends of asymptomatic parasitaemia in a seasonal transmission setting in North-Western Burkina Faso between 2000 and 2009-2012. Malar J 12, 27 (2013).

7. Adjah J, Fiadzoe B, Ayanful-Torgby R, Amoah LE. Seasonal variations in Plasmodium falciparum genetic diversity and multiplicity of infection in asymptomatic children living in southern Ghana. BMC Infect Dis 18, 432 (2018).

8. Ippolito MM, et al. The Relative Effects of Artemether-lumefantrine and Non-artemisinin Antimalarials on Gametocyte Carriage and Transmission of Plasmodium falciparum: A Systematic Review and Meta-analysis. Clin Infect Dis 65, 486–494 (2017).

9. Eksi S, et al. Plasmodium falciparum gametocyte development 1 (Pfgdv1) and gametocytogenesis early gene identification and commitment to sexual development. PLoS Pathog 8, e1002964 (2012).

10. Kafsack BF, et al. A transcriptional switch underlies commitment to sexual development in malaria parasites. Nature 507, 248–252 (2014).

11. Filarsky M, et al. GDV1 induces sexual commitment of malaria parasites by antagonizing HP1-dependent gene silencing. Science 359, 1259–1263 (2018).

12. Josling GA, et al. Dissecting the role of PfAP2-G in malaria gametocytogenesis. Nat Commun 11, 1503 (2020).

13. Salcedo-Amaya AM, et al. Dynamic histone H3 epigenome marking during the intraerythrocytic cycle of Plasmodium falciparum. Proc Natl Acad Sci U S A 106, 9655–9660 (2009).

14. Flueck C, et al. Plasmodium falciparum heterochromatin protein 1 marks genomic loci linked to phenotypic variation of exported virulence factors. PLoS Pathog 5, e1000569 (2009).

15. Sales-Gil R, Vagnarelli P. How HP1 Post-Translational Modifications Regulate Heterochromatin Formation and Maintenance. Cells 9, (2020).

16. Papamokos GV, Kaxiras E. How to evict HP1 from H3: Using a complex salt bridge. Biophys Chem 300, 107062 (2023).

17. Hirota T, Lipp JJ, Toh BH, Peters JM. Histone H3 serine 10 phosphorylation by Aurora B causes HP1 dissociation from heterochromatin. Nature 438, 1176–1180 (2005).

18. Fischle W, et al. Regulation of HP1-chromatin binding by histone H3 methylation and phosphorylation. Nature 438, 1116–1122 (2005).

19. Treeck M, Sanders JL, Elias JE, Boothroyd JC. The phosphoproteomes of Plasmodium falciparum and Toxoplasma gondii reveal unusual adaptations within and beyond the parasites’ boundaries. Cell Host Microbe 10, 410–419 (2011).

20. Ganter M, et al. Plasmodium falciparum CRK4 directs continuous rounds of DNA replication during schizogony. Nat Microbiol 2, 17017 (2017).

21. Pease BN, et al. Global analysis of protein expression and phosphorylation of three stages of Plasmodium falciparum intraerythrocytic development. J Proteome Res 12, 4028–4045 (2013).

22. Amos B, et al. VEuPathDB: the eukaryotic pathogen, vector and host bioinformatics resource center. Nucleic Acids Res 50, D898–D911 (2022).

23. Zhang M, et al. Uncovering the essential genes of the human malaria parasite Plasmodium falciparum by saturation mutagenesis. Science 360, (2018).

24. Tiburcio M, et al. A 39-Amino-Acid C-Terminal Truncation of GDV1 Disrupts Sexual Commitment in Plasmodium falciparum. mSphere 6, (2021).

25. Llora-Batlle O, et al. Conditional expression of PfAP2-G for controlled massive sexual conversion in Plasmodium falciparum. Sci Adv 6, eaaz5057 (2020).

26. MalariaGen, et al. Pf7: an open dataset of Plasmodium falciparum genome variation in 20,000 worldwide samples. Wellcome Open Res 8, 22 (2023).

27. Donnelly MLL, et al. The ‘cleavage’ activities of foot-and-mouth disease virus 2A site-directed mutants and naturally occurring ‘2A-like’ sequences. J Gen Virol 82, 1027–1041 (2001).

28. Liu Z, et al. Systematic comparison of 2A peptides for cloning multi-genes in a polycistronic vector. Sci Rep 7, 2193 (2017).

29. Santos JM, et al. Red Blood Cell Invasion by the Malaria Parasite Is Coordinated by the PfAP2-I Transcription Factor. Cell Host Microbe 21, 731–741 e710 (2017).

30. Shang X, et al. A cascade of transcriptional repression determines sexual commitment and development in Plasmodium falciparum. Nucleic Acids Res 49, 9264–9279 (2021).

31. Subudhi AK, et al. DNA-binding protein PfAP2-P regulates parasite pathogenesis during malaria parasite blood stages. Nat Microbiol 8, 2154–2169 (2023).

32. Yuda M, Kaneko I, Murata Y, Iwanaga S, Nishi T. Mechanisms of triggering malaria gametocytogenesis by AP2-G. Parasitol Int 84, 102403 (2021).

33. Kent RS, Modrzynska KK, Cameron R, Philip N, Billker O, Waters AP. Inducible developmental reprogramming redefines commitment to sexual development in the malaria parasite Plasmodium berghei. Nat Microbiol 3, 1206–1213 (2018).

34. Kato N, et al. Diversity-oriented synthesis yields novel multistage antimalarial inhibitors. Nature 538, 344–349 (2016).

35. Baniecki ML, Wirth DF, Clardy J. High-throughput Plasmodium falciparum growth assay for malaria drug discovery. Antimicrob Agents Chemother 51, 716–723 (2007).

36. Jillapalli R, et al. A Plasmodium berghei putative serine-threonine kinase 2 (PBANKA_0311400) is required for late liver stage development and timely initiation of blood stage infection. Biol Open 8, (2019).

37. Ikadai H, et al. Transposon mutagenesis identifies genes essential for Plasmodium falciparum gametocytogenesis. Proc Natl Acad Sci U S A 110, E1676–1684 (2013).

38. Nguyen W, et al. 7-N-Substituted-3-oxadiazole Quinolones with Potent Antimalarial Activity Target the Cytochrome bc(1) Complex. ACS Infect Dis 9, 668–691 (2023).

39. Antonova-Koch Y, et al. Open-source discovery of chemical leads for next-generation chemoprotective antimalarials. Science 362, (2018).

40. Hayward JA, et al. A screen of drug-like molecules identifies chemically diverse electron transport chain inhibitors in apicomplexan parasites. PLoS Pathog 19, e1011517 (2023).

41. Calit J, et al. Pyrimidine Azepine Targets the Plasmodium bc (1) Complex and Displays Multistage Antimalarial Activity. JACS Au 4, 3942–3952 (2024).

42. Prajapati SK, Joshi H, Dev V, Dua VK. Molecular epidemiology of Plasmodium vivax anti-folate resistance in India. Malar J 10, 102 (2011).

43. Beckmann JF, Van Vaerenberghe K, Akwa DE, Cooper BS. A single mutation weakens symbiont-induced reproductive manipulation through reductions in deubiquitylation efficiency. Proc Natl Acad Sci U S A 118, (2021).

44. Kazemi-Esfarjani P, et al. Substitution of valine-865 by methionine or leucine in the human androgen receptor causes complete or partial androgen insensitivity, respectively with distinct androgen receptor phenotypes. Mol Endocrinol 7, 37–46 (1993).

45. Walss-Bass C, et al. A novel missense mutation in the transmembrane domain of neuregulin 1 is associated with schizophrenia. Biol Psychiatry 60, 548–553 (2006).

46. Rajebhosale P, et al. Neuregulin1 nuclear signaling influences adult neurogenesis and regulates a schizophrenia susceptibility gene network within the mouse dentate gyrus. bioRxiv, 2022.2008.2010.503469 (2024).

47. Adjalley S, Lee MCS. CRISPR/Cas9 Editing of the Plasmodium falciparum Genome. Methods Mol Biol 2470, 221–239 (2022).

48. Ghorbal M, Gorman M, Macpherson CR, Martins RM, Scherf A, Lopez-Rubio JJ. Genome editing in the human malaria parasite Plasmodium falciparum using the CRISPR-Cas9 system. Nat Biotechnol 32, 819–821 (2014).

49. Heigwer F, Kerr G, Boutros M. E-CRISP: fast CRISPR target site identification. Nat Methods 11, 122–123 (2014).

50. Untergasser A, et al. Primer3--new capabilities and interfaces. Nucleic Acids Res 40, e115 (2012).

51. Li H, Durbin R. Fast and accurate short read alignment with Burrows-Wheeler transform. Bioinformatics 25, 1754–1760 (2009).

52. DePristo MA, et al. A framework for variation discovery and genotyping using next-generation DNA sequencing data. Nat Genet 43, 491–498 (2011).

53. https://www.genome.jp/tools/motif/.

54. Campbell TL, De Silva EK, Olszewski KL, Elemento O, Llinas M. Identification and genome-wide prediction of DNA binding specificities for the ApiAP2 family of regulators from the malaria parasite. PLoS Pathog 6, e1001165 (2010).

55. One-way ANOVA followed by Dunnett’s multiple comparisons test was performed using GraphPad Prism version 10.0.0 for Windows, GraphPad Software, Boston, Massachusetts USA, www.graphpad.com.

56. Bindels DS, et al. mScarlet: a bright monomeric red fluorescent protein for cellular imaging. Nat Methods 14, 53–56 (2017).

57. Anna O. Chertkova MM, Marten Postma, Nikki van Bommel, Sanne van der Niet, Kevin L. Batenburg, Linda Joosen, Theodorus W.J. Gadella Jr., Yasushi Okada, Joachim Goedhart. Robust and Bright Genetically Encoded Fluorescent Markers for Highlighting Structures and Compartments in Mammalian Cells. bioRxiv (2017).

58. Zhang W, et al. Optogenetic control with a photocleavable protein, PhoCl. Nat Methods 14, 391–394 (2017).

59. Sinha A, et al. A cascade of DNA-binding proteins for sexual commitment and development in Plasmodium. Nature 507, 253–257 (2014).

