## Supplementary figures and tables for "A single valine to leucine switch disrupts *Plasmodium falciparum* AP2-G DNA binding and reveals GDV1’s role in *ap2-g* activation"

[1]Department of Microbiology and Immunology, Uniformed Services University of the Health Sciences, Bethesda, USA, [2]Henry M. Jackson Foundation for the Advancement of Military Medicine, Inc., Bethesda, USA, [3]Laboratory of Malaria and Vector Research, NIAID, NIH, Rockville, USA, [4]Department of Molecular Biology, Lewis-Sigler Institute for Integrative Genomics, Princeton University, Princeton, USA, [5]Department of Anatomy, Physiology & Genetics, Uniformed Services University of the Health Sciences, Bethesda, USA, [6]Department of Microbiology & Immunology, Weill Cornell Medicine, New York, USA, [7]Department of Biochemistry & Molecular Biology and the Huck Center for Malaria Research, Pennsylvania State University, University Park, USA, [8]Department of Chemistry, Pennsylvania State University, University Park, USA. *Corresponding author, [@]present address: Integrative Genomics and Bioinformatics Core, The Salk Institute for Biological Studies, La Jolla, CA, USA, [#]current affiliation.

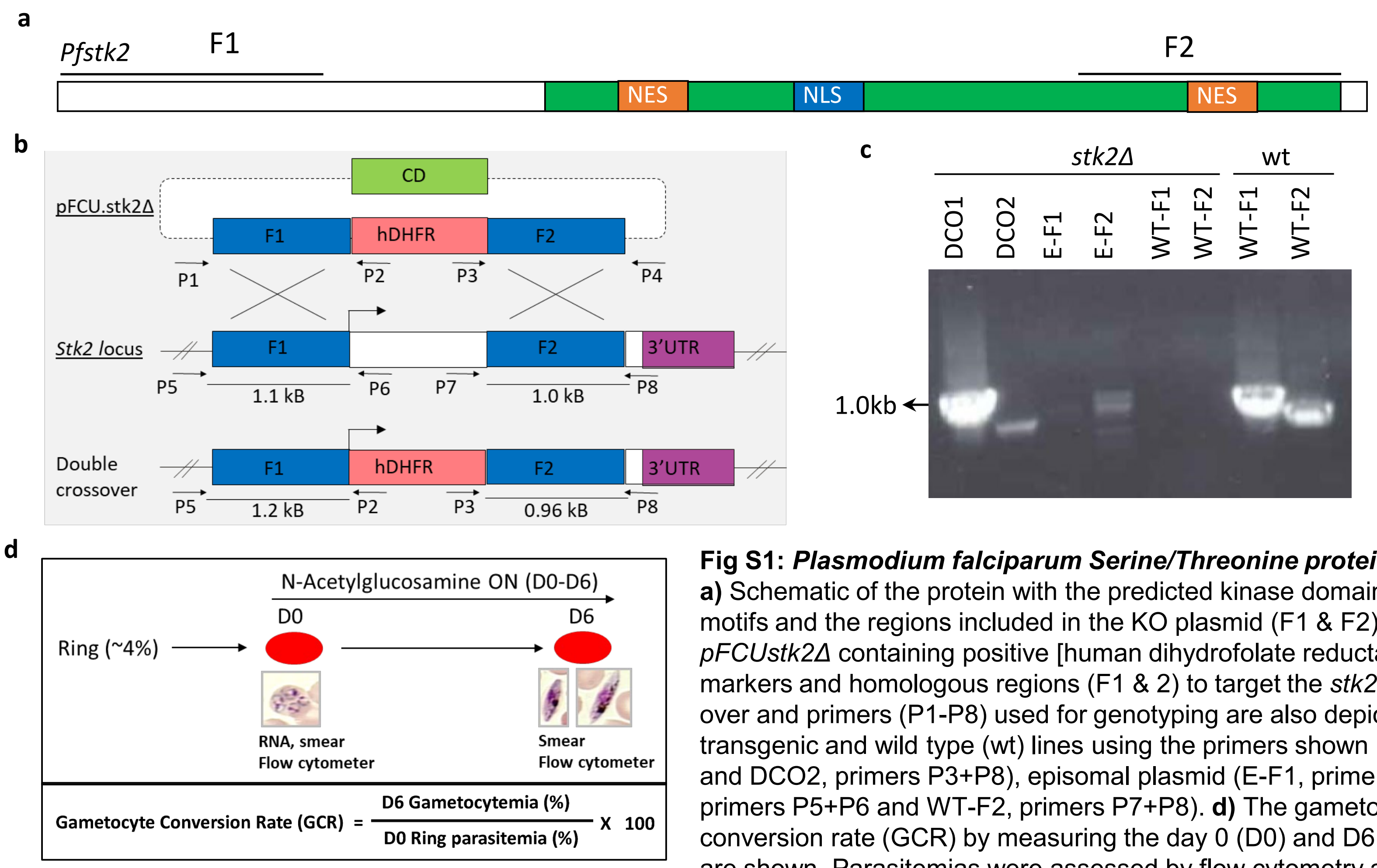


**Fig S1:** ***Plasmodium falciparum* Serine/Threonine protein kinase** (*Pfstk2*) knockout strategy and gametocyte phenotyping assay. **a)** Schematic of the protein with the predicted kinase domain (green), nuclear export signal (NES), nuclear localization signal (NLS) motifs and the regions included in the KO plasmid (F1 & F2) indicated**. B)** The double crossover *knockout* strategy using plasmid *pFCUstk2Δ* containing positive [human dihydrofolate reductase (DHFR)] and negative [Cytosine deaminase (CD)] selectable markers and homologous regions (F1 & 2) to target the *stk2* is shown. The *Pfstk2* locus before and after a successful double cross over and primers (P1-P8) used for genotyping are also depicted. **c)** Polymerase chain reaction (PCR) genotyping of the *3D7.stk2Δ* transgenic and wild type (wt) lines using the primers shown in **b** to detect double crossover (DCO) integration (DCO1, primers P2+P5 and DCO2, primers P3+P8), episomal plasmid (E-F1, primers P1+P2 and E-F2, primers P3+P4) and the wild type locus (WT-F1, primers P5+P6 and WT-F2, primers P7+P8). **d)** The gametocyte phenotyping assay and formula used to determine the gametocyte conversion rate (GCR) by measuring the day 0 (D0) and D6 parasitemias of cultures grown in the presence of N-acetylglucosamine are shown. Parasitemias were assessed by flow cytometry and Giemsa-stained culture smears.

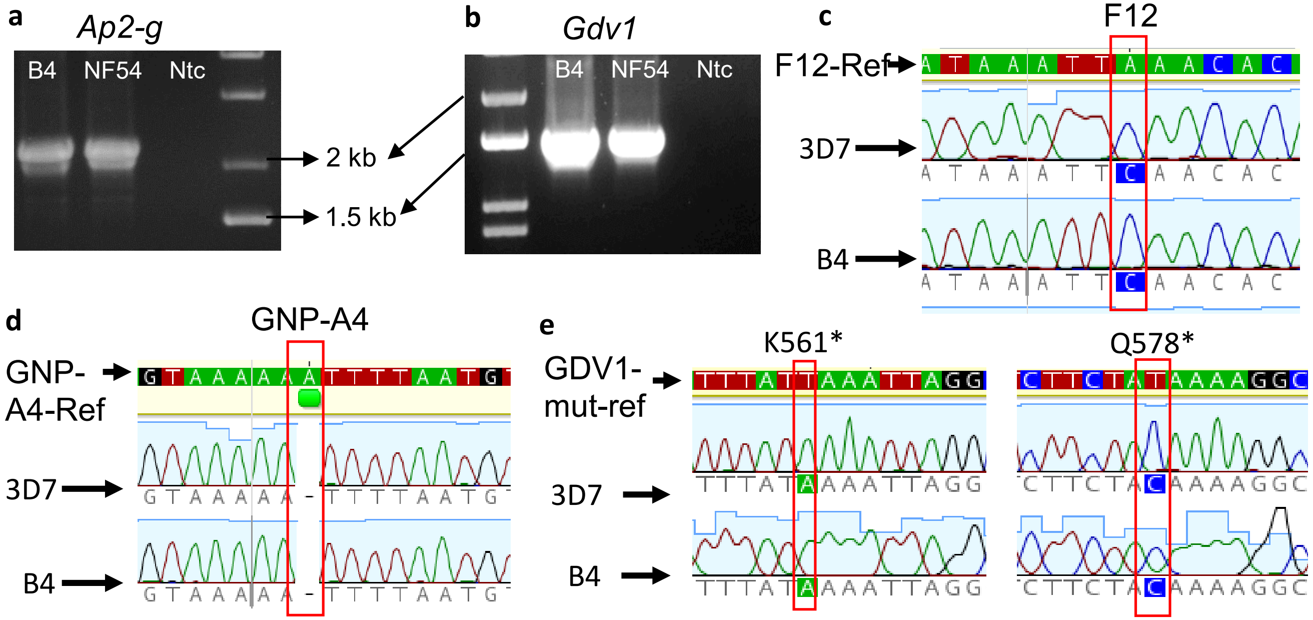


**Fig S2: *Ap2-g* and *gdv1* loci test for known mutations conferring premature termination of protein.** *Ap2-g* from coding region (bp 6024) to 3'-UTR (bp 988) **(a)** and *gdv1* from coding region (bp 841) to 3'-UTR (bp 532) **(b)** were amplified using polymerase chain reaction from *3D7.stk2Δ* clone B4, or wild type (wt) parasite DNA and a no template control (Ntc). **c-e)** Chromatograms of the DNA sequence of wt 3D7 or *3D7.stk2Δ clone B4* (B4) parasites are shown below the reference sequence for the known *ap2-g* [1] **(c,d)** or *gdv1* [2] **(e)** [3] **(f)** mutations that block gametocyte production. Red boxes (**c-e**) indicates the nucleotide positions where mutations in *ap2-g* were expected for parasite lines F12 and GNP-A4 and in *gdv1* (K561*, Q578*).

**Reference:**

1) Kafsack BF, et al. A transcriptional switch underlies commitment to sexual development in malaria parasites. Nature 507, 248-252 (2014).
2) Tiburcio M, et al. A 39-Amino-Acid C-Terminal Truncation of GDV1 Disrupts Sexual Commitment in Plasmodium falciparum. mSphere 6, (2021).
3) Llora-Batlle O, et al. Conditional expression of PfAP2-G for controlled massive sexual conversion in Plasmodium falciparum. Sci Adv 6, eaaz5057 (2020).

| 2163th ↓ | AP2-G (2159-2218) | Pridicted AP2 domain Position | i-Evalue |
|---|---|---|---|
| Wt (V) | PIHSVWKDTTRGHCSWRCRWWENGRRLSKNFNVKRFGNDGALRMAITMKLKKSNPKEQMQ | 2162 - 2210 | 0.00053 |
| V->M | PIHSMWKDTTRGHCSWRCRWWENGRRLSKNFNVKRFGNDGALRMAITMKLKKSNPKEQMQ | 2173 - 2210 | 0.0023 |
| V->V | PIHSFWKDTTRGHCSWRCRWWENGRRLSKNFNVKRFGNDGALRMAITMKLKKSNPKEQMQ | 2173 - 2210 | 0.0023 |
| V->W | PIHSWWKDTTRGHCSWRCRWWENGRRLSKNFNVKRFGNDGALRMAITMKLKKSNPKEQMQ | 2173 - 2210 | 0.0023 |
| V->Y | PIHSYWKDTTRGHCSWRCRWWENGRRLSKNFNVKRFGNDGALRMAITMKLKKSNPKEQMQ | 2173 - 2210 | 0.0023 |
| V->S | PIHSSWKDTTRGHCSWRCRWWENGRRLSKNFNVKRFGNDGALRMAITMKLKKSNPKEQMQ | 2173 - 2210 | 0.0023 |
| V->T | PIHSTWKDTTRGHCSWRCRWWENGRRLSKNFNVKRFGNDGALRMAITMKLKKSNPKEQMQ | 2173 - 2210 | 0.0023 |
| V->N | PIHSNWKDTTRGHCSWRCRWWENGRRLSKNFNVKRFGNDGALRMAITMKLKKSNPKEQMQ | 2173 - 2210 | 0.0023 |
| V->Q | PIHSQWKDTTRGHCSWRCRWWENGRRLSKNFNVKRFGNDGALRMAITMKLKKSNPKEQMQ | 2173 - 2210 | 0.0023 |
| V->C | PIHSCWKDTTRGHCSWRCRWWENGRRLSKNFNVKRFGNDGALRMAITMKLKKSNPKEQMQ | 2173 - 2210 | 0.0023 |
| V->P | PIHSPWKDTTRGHCSWRCRWWENGRRLSKNFNVKRFGNDGALRMAITMKLKKSNPKEQMQ | 2173 - 2210 | 0.0023 |
| V->R | PIHSRWKDTTRGHCSWRCRWWENGRRLSKNFNVKRFGNDGALRMAITMKLKKSNPKEQMQ | 2173 - 2210 | 0.0023 |
| V->H | PIHSHWKDTTRGHCSWRCRWWENGRRLSKNFNVKRFGNDGALRMAITMKLKKSNPKEQMQ | 2173 - 2210 | 0.0023 |
| V->K | PIHSKWKDTTRGHCSWRCRWWENGRRLSKNFNVKRFGNDGALRMAITMKLKKSNPKEQMQ | 2173 - 2210 | 0.0023 |
| V->D | PIHSDWKDTTRGHCSWRCRWWENGRRLSKNFNVKRFGNDGALRMAITMKLKKSNPKEQMQ | 2173 - 2210 | 0.0023 |
| V->E | PIHSEWKDTTRGHCSWRCRWWENGRRLSKNFNVKRFGNDGALRMAITMKLKKSNPKEQMQ | 2173 - 2210 | 0.0023 |

**Fig S3: *In-silico* replacement of valine with other amino acids in AP2-G AP2 domain predictions using MOTIF search software.** AP2-G amino acids (aa) 2159-2218 are shown from wild type 3D7 and with valine$_{2163}$ substituted with the indicated amino acid. Red text indicates the AP2 domain predicted for each sequence using MOTIF search software and the green box indicates position 2163. The last two columns are the aa included in the predicted AP2 domain and the i-Evalue of the sequence that predicts the DNA interaction strength.

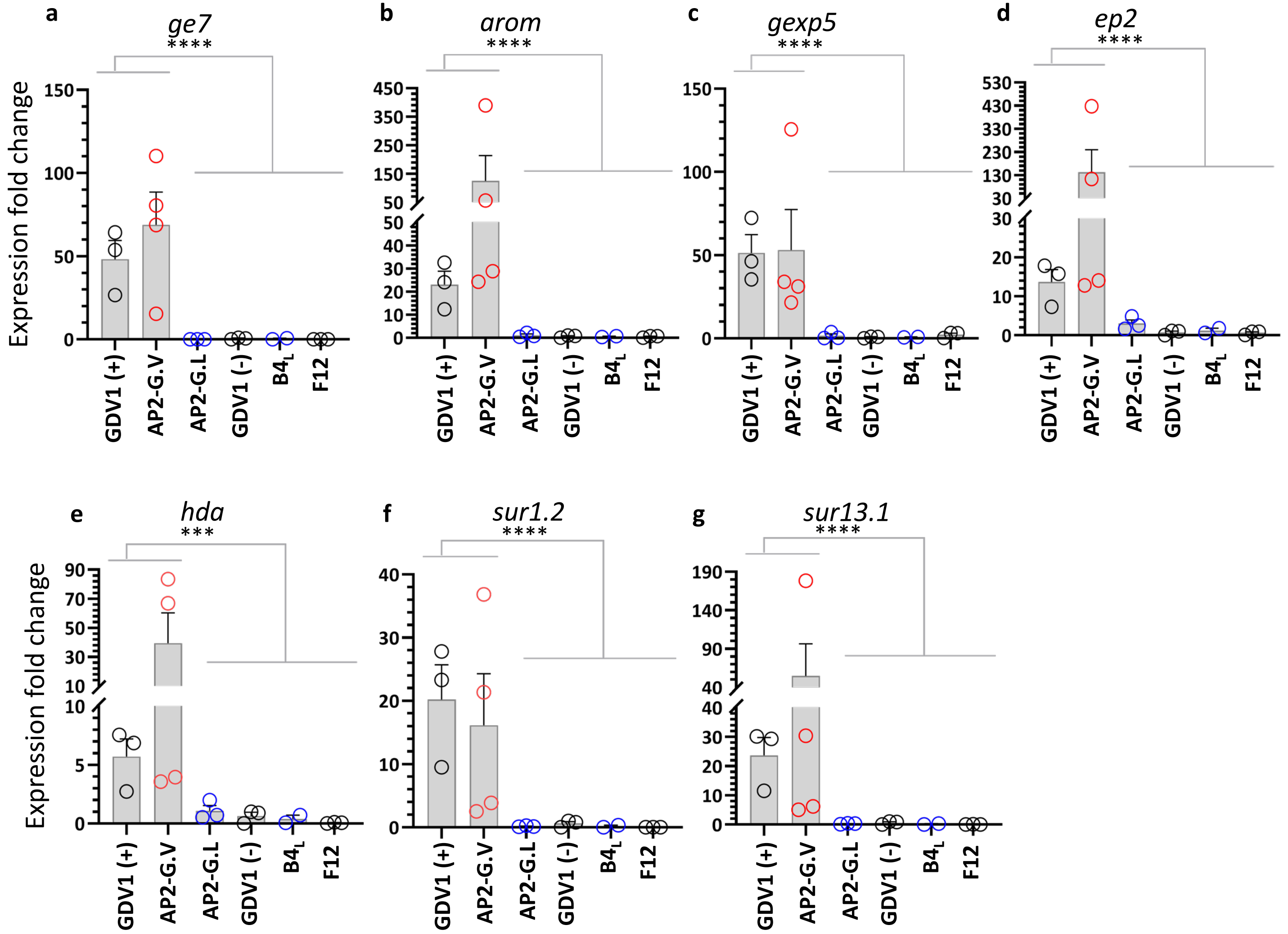


**Fig S4: Ring stage expression of AP2-G dependent genes**. AP2-G dependent gene expression was analyzed using reverse transcriptase–quantitative polymerase chain reaction (RT-qPCR) of RNA isolated from sorbitol synchronized ring stage parasites (6-10 hpi) encoding *ap2-g.V [NF54.gdv1.3xha.fkbp* with (GDV1(+) or without (GDV1(-)) Shld1, *3D7.stk2Δ//ap2-g.V* (AP2-G-V)], *ap2-g. L*, [*NF54.ap2-g.L* (AP2-G.L), *3D7.stk2Δ//ap2-g.L* ($B4_L$)] or truncated *ap2-G* (F12). The GDV1 (-) parasite line was used as the comparator to determine fold change. The RT-qPCR primers used are listed in **Supplemental Table 2** with the PlasmoDB ID # of the gene. Each circle indicates an independent data point and the bar and error bar represent the mean and standard error of the mean, respectively. A Kruskal-Wallis test followed by Dunn's multiple comparison was used for statistical analysis between groups. The p value of each comparison is indicated by *** (<0.001) and **** (<0.0001).

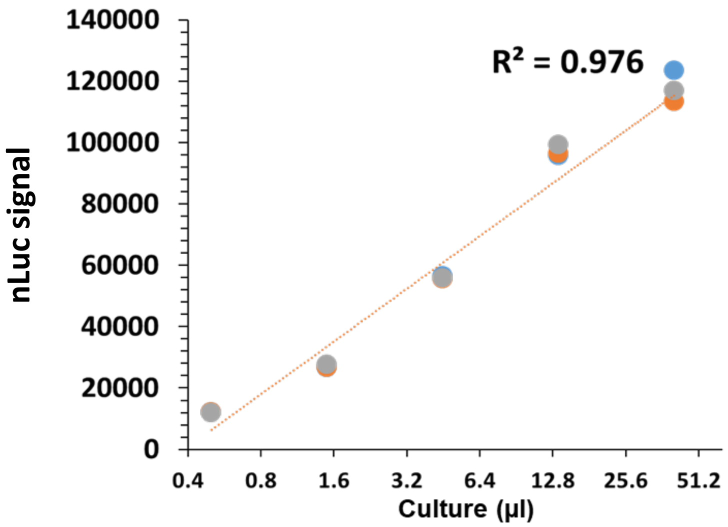


**Fig S5. Nanoluciferase (nLuc) signal is positively correlated with parasite number (culture volume) using an *ap2-g* reporter line, *NF54.ap2-g.V.p2a.nLuc*.** Colored circles indicate biological replicates. All circles are shown for the culture volumes, but for each of the first 3 volumes the points overlap.

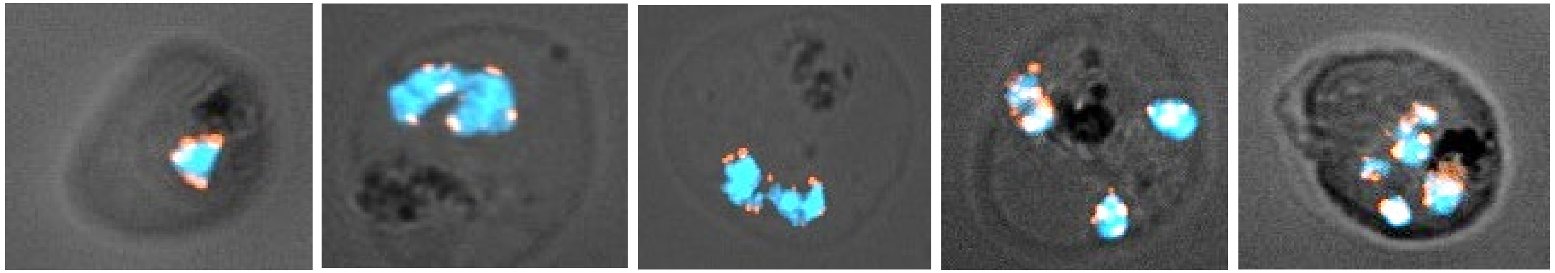

**Fig S6. GDV1 expression during schizogony begins in late trophozoites and continues into early schizonts.** *683.Gdv1.fkbp.tdTom* transgenic parasites were stained with Hoechst (1:500 dilution) and imaged using fluorescence microscopy. An overlay of the bright field, Hoechst and tdTomato images are shown.

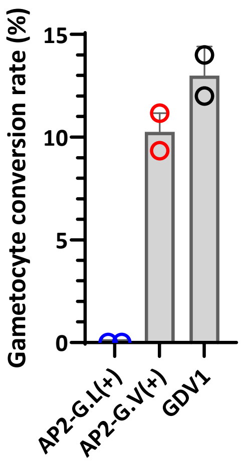


**Fig. S7: Day 6 gametocyte conversion rate of the parasite lines used in the nanoluciferase (nLuc) time course (Figure 5a).** Day 6 gametocyte conversion rates of AP2-G.L(+), *3D7.stk2Δ//ap2-g.L.fkbp.p2a.nLuc* (Shld1+,blue circle), AP2-G.V (+), *3D7.stk2Δ //ap2g.V.fkbp.p2a.nLuc* (Shld1+,red circle), and GDV1, *NF54.gdv1.p2a.nLuc* (black circle) are shown. Each circle indicates an independent data point and the bar and error bars represent mean and standard error of the mean, respectively.

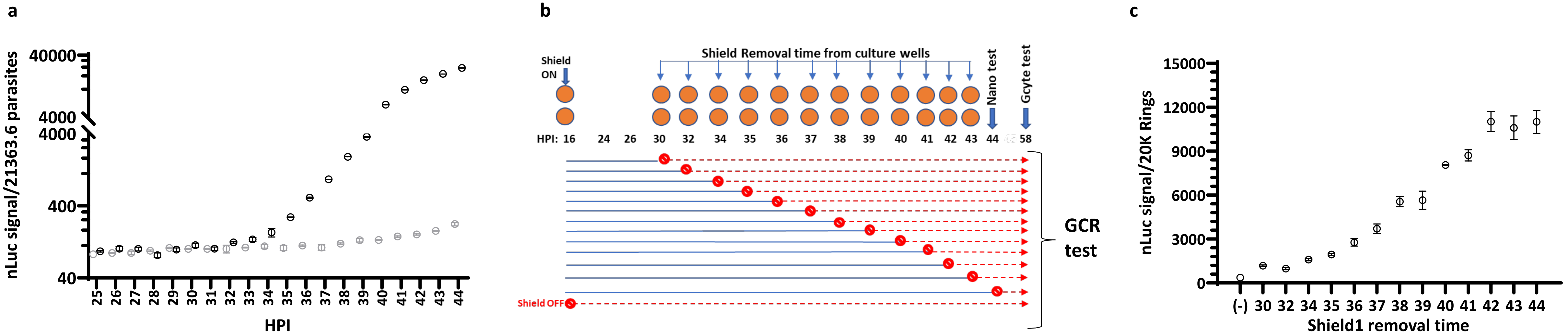


**Fig. S8: Time-course to determine when GDV1 is required for maximal *ap2-g* expression. a)** Time-course of *ap2-g*-linked nanoluciferase (nLuc) activity using the *NF54.gdv1.3xha.fkbp//ap2-g.p2 nLuc* line in the presence (black) and absence (gray) of Shld1. **b)** Schematic showing the time points when GDV1 was first added and then removed from the culture wells. Blue horizontal lines indicate culture wells with Shld1 whereas red dotted lines indicate no Shld1. At 58 hpi, after monitoring nLuc, N-acetylglucosamine (NAG) was added to each well and this was designated Day 0 for gametocyte assessment and gametocytemia was measured 7 days later (D6) to calculate gametocyte conversion rate (GCR). **c)** The 56 hpi *ap2-g-* linked nLuc signal of each well before adding NAG. Each circle indicates an independent data point and the error bars represent the standard error of the mean, respectively.

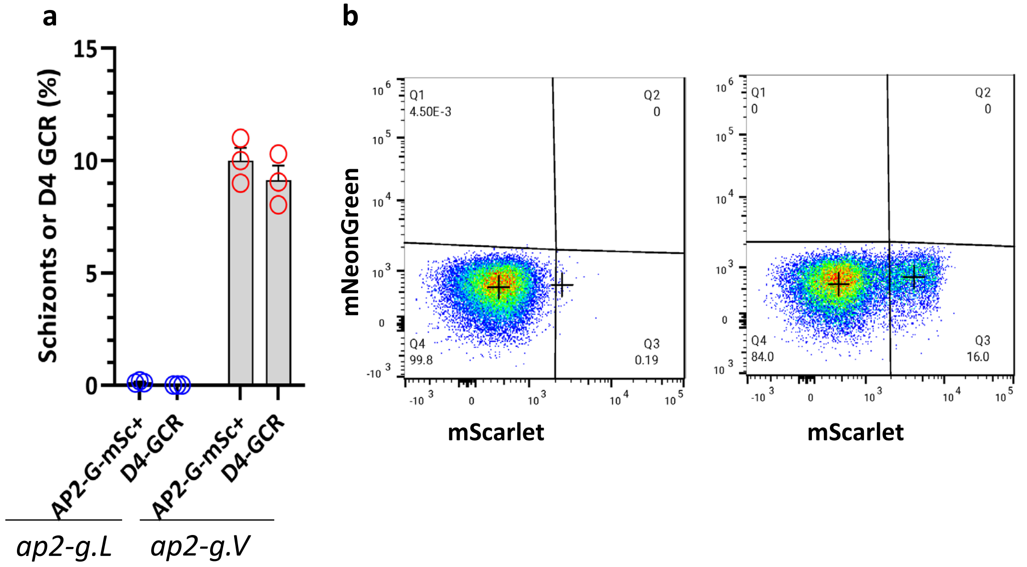


**Fig. S9: Flow cytometric analysis of AP2.V and AP2.L reporter lines.**
**a)** Comparison of the % of AP2-G.mSc+ schizonts detected by flow cytometry and the day 4 gametocyte conversion rate (D4-GCR) of *ap2-g.L* (*3D7.stk2Δ//ap2-g.L.mSc*) and *ap2-g.V* (*3D7.stk2Δ//ap2-g.V.mSc*) mScarlet reporter lines. **b)** Flow cytometric analysis of *ap2-g.L* (*3D7.stk2Δ//ap2-g.L.mSc*) (left panel) and *ap2-g.V* (*3D7.stk2Δ//ap2-g.V.mSc*) (right panel) mScarlet reporter lines.

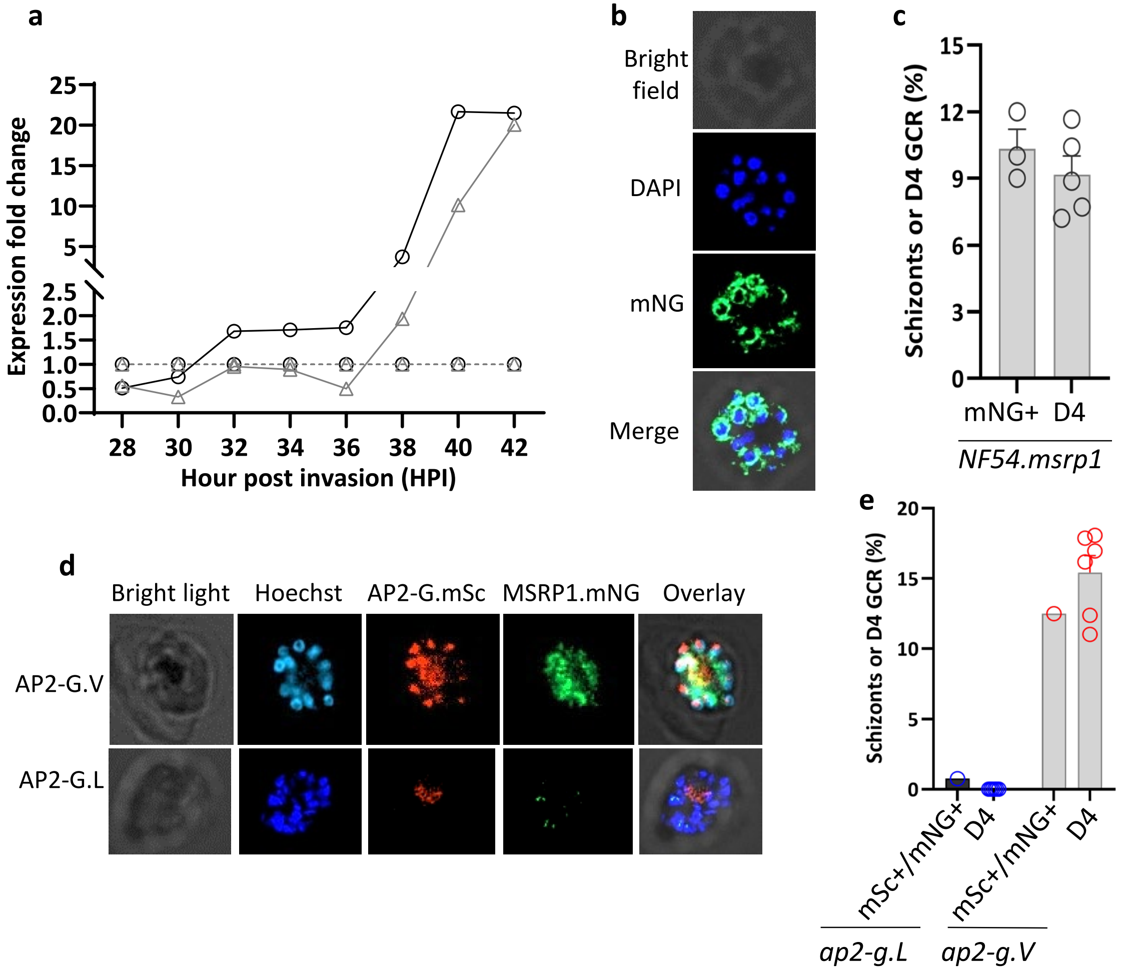


**Fig S10: *Msrp1* and *ap2-g* expression time-course during schizogony**

**a)** The increase in *msrp1* (triangles) and *ap2-g* (circles) RNA over time in a *gdv1* inducible line *NF54.gdv1.3xha.fkbp* in the presence of Shld1 (continuous line) or absence of Shld1 (dotted line). **b**) Live imaging of the *NF54.msrp1.mNG* line stained with Hoechst to detect DNA demonstrates MSRP1.mNG expression in mature schizonts. **c)** Comparison of the % of MSRP1.mNG positive schizonts measured by flow cytometry and the day 4 gametocyte conversion rate in *NF54.msrp1.mNG* parasites. **d)** Live imaging of AP2-G.mSc and MSRP1.mNG expression in the *NF54.msrp1.mNG//ap2-g.V.mSc* (top panel) and *NF54.msrp1.mNG//ap2-g.L.mSc* (bottom panel) lines stained with Hoechst to detect DNA. **e)** Comparison of the % of double positive schizonts (AP2-G.mSc+/MSRP1.mNG+) and the day 4 gametocyte conversion rate in *ap2-L*, *3D7.stk2Δ//ap2-g.L.mSc//msrp1.mNG*) or *ap2-g.V* (*3D7.stk2Δ//ap2-g.V.mSc//msrp1.mNG*) expressing lines. Blue and red circles indicate *ap2-g.L* and *ap2-g.V* lines, respectively. Each circle indicates independent data points and the bar and error bars represent the mean and standard error of the mean, respectively.

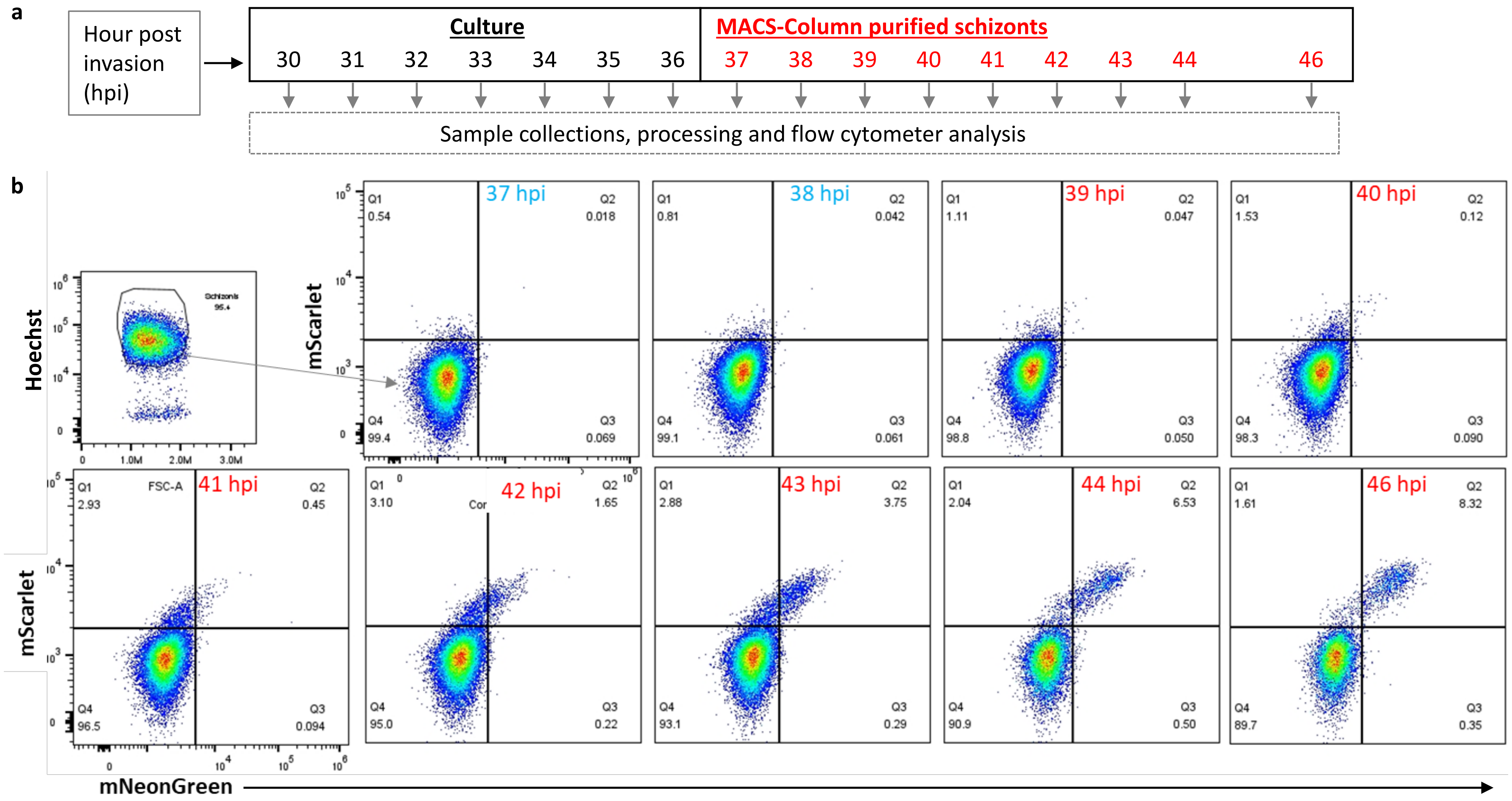


**Fig. S11: Time-course of AP2-G.mScarlet and MSRP1.mNeonGreen expression through schizogony. a)** Schematic of the experimental design. Beginning 30 h post RBC invasion (hpi) samples were taken from a synchronized culture for flow cytometric analysis. At 36 hpi schizonts had matured enough to allow MACS-column purification before flow cytometry, which reduced the number of uninfected RBCs. **b**) Quantification of AP2-G.mSc and MSRP1.mNG positive schizonts using flow cytometry. None of the early schizonts were positive for AP2-G.mSc before 37 hpi and therefore not included in the figure. AP2-G.mSc positive schizonts gradually increased from 38 hpi reaching 1% of the schizont population at 39 hpi. The % of AP2-G.mSc single positive schizonts plateaued at ~3% at 41 hr as MSRP1.mNG expression was detected and the number of double positive schizonts began to increase. After 43 hr the number of AP2-G single positive schizonts declined rapidly, while the number of double positive schizonts continued to increase, again consistent with AP2-G expression being required for MSRP1 expression.

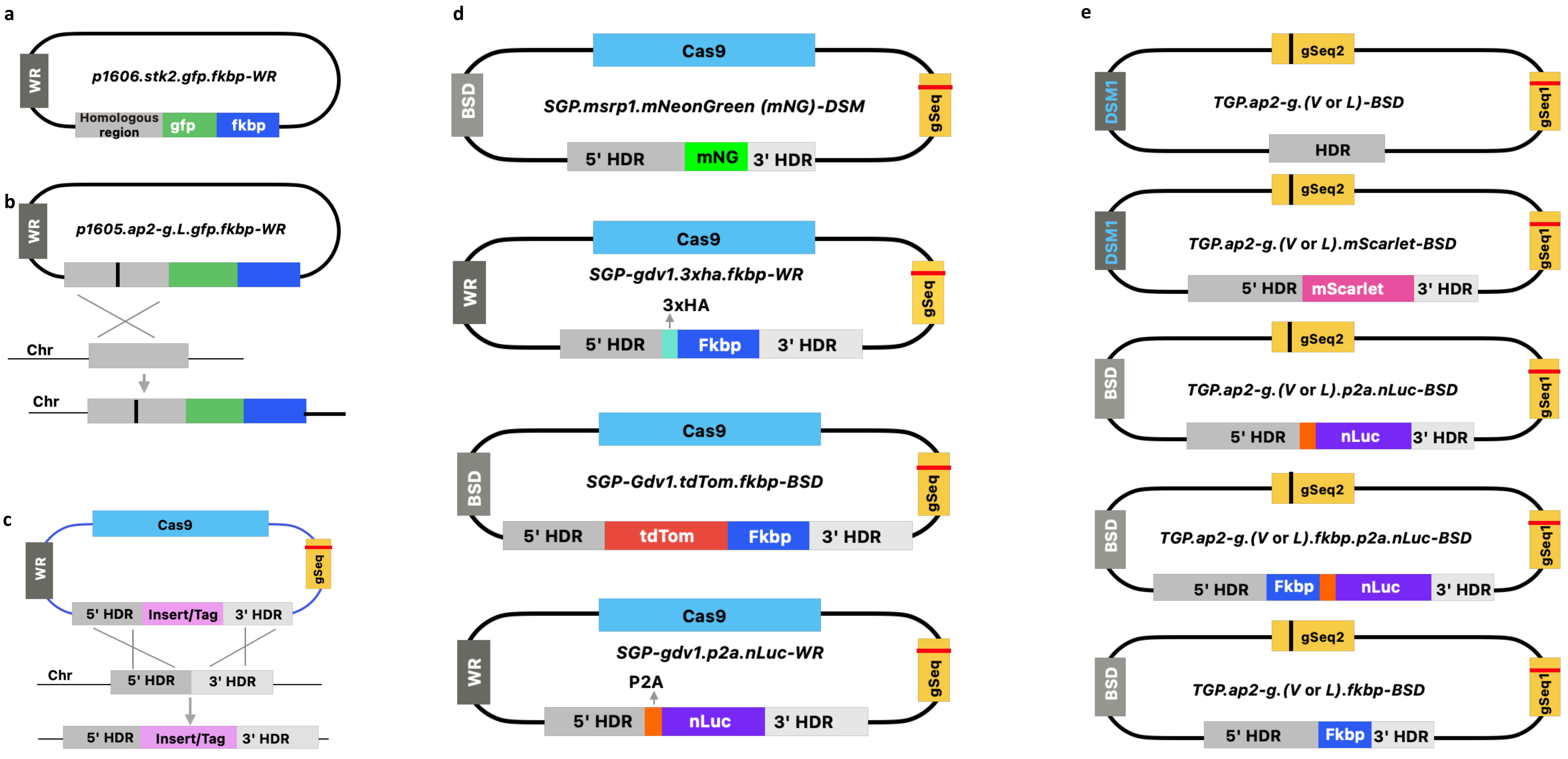


**Fig. S12: Plasmids used to tag/modify *stk2, gdv1, ap2-g* and *msrp1*.** The selectable markers were *dihydrofolate reductase* (WR drug), *blasticidin S deaminase* (BSD drug), *dihydroorotate dehydrogenase* (DSM1 drug). Black line in the homologous region **(b)** indicates *ap2-g.L* mutation. HDR and gSeq **(d-e)** indicated homology directed repair template and guide RNA sequence, respectively. Fkbp is an inducible expression system that targets the attached protein for degradation in the absence of Shld1 ligand.

**Table S1: Whole genome sequences showing mutations in B4 clone (*3D7.stk2Δ//ap2-g.L* ) and wild type NF54.**

| Chromosome | Position | Annotation | Gene | Ref Allele | Alt allele | GT:AD:DP (B4 clone) | GT:AD:DP (NF54) |
|---|---|---|---|---|---|---|---|
| Pf3D7_10_v3 | 1187187 | missense | PF3D7_1029000 | G | A | 1/1:2,27:30 | 1/1:4,26:34 |
| Pf3D7_10_v3 | 1437030 | missense | PF3D7_1036400 | A | G | 1/1:0,8:8 | ./. (Deletion) |
| Pf3D7_12_v3 | 913689 | missense | PF3D7_1222600 | **G** | **T** | 1/1:0,101:101 | 0/0:.:143 |
| Pf3D7_12_v3 | 1908160 | missense | PF3D7_1245800 | A | T | 1/1:0,17:17 | 1/1:0,20:20 |
| Pf3D7_12_v3 | 1908195 | missense | PF3D7_1245800 | T | A | 1/1:0,32:32 | 1/1:0,28:28 |

GT: Genotype, AD: Read depth for each allele, DP: Read depth

**Table S2: List of primers used for plasmid construction, transgenic line genotyping and genes expression analysis.**

| Oligo types | Oligo name | Oligo sequence (5' - 3') (Ref) | Ref |
|---|---|---|---|
| **Plasmid construction** | | | |
| *Stk2*-KO (PF3D7_0214600) | *P21 (P1)* | *TCCCCGCGGTTAAGTACAATTACATGTAATTAA* | |
| | *P22 (P2)* | *CTAGACTAGTGTTCTTATGATTAGTAATAATTTC* | |
| | *P68 (P3)* | *CATGCCATGGTTGAAGAGCCCAAAATGTTAT* | |
| | *P69 (P4)* | *GATCCTAGGCAAATCTCTAAAAGAATAATG* | |
| *Stk2 (PF3D7_0214600)* | *STK2(Xho1)_F* | *TAATCTCGAGTATTCAATTGAAGAGCCC* | |
| | *STK2(Xho1)_R* | *TAATCCTAGGTATCAAATCTCTAAAAGAATAATG* | |
| *Gdv1 (PF3D7_0935400)* | GDV1_recoGseq1_F | tattGGATTTATTCTCATAACAAC | |
| | GDV1_recoGseq1_R | aaacGTTGTTATGAGAATAAATCC | |
| | GDV1_InFu_F | GTACCGAGCTCGAATTCTTGATAAAATATGGGAGATC | |
| | GDV1_recoHDR1_R | GGGTTGATACGCATTACGACTGGAATACCTTTCTGTTTTA | |
| | GDV1_recoHDR1_F | AGTCGTAATGCGTATCAACCCTTTTCAATCTTTTCGTTTA | |
| | GDV1_InFu_Rn | CATATGGGTAGCGGCCGCTTTTATATGTACATTTTTC | |
| | GDV1 3UTR_InFu_F | AACTAGTGTGACCGGTTAAATAAATGAAAAAAAAAA | |
| | GDV1 3UTR_InFu_R | AGTGCCACCTGACGTCATGTTTTTATTTGTGCTTA | |
| | GDV1_InFu_tdTom_F | CATATAAAAGCGGCCGCATGGTGAGCAAGGGCGAGG | |
| | GDV1_InFu_FKBP_R | CATTTATTTAACCGGTTTCTTCCGGTTTTAGAAGC | |
| | Gdv1_P2ANano_F | ACATATAAAAGCGGCCGCGCCACAAACTTCTCTCTGCT | |
| | Gdv1_Nano_R | TTAACCGGTCACACTAGTCGCCAGAATGCGTTCGCACAG | |
| *Ap2-g (PF3D7_1222600)* | *AP2-g Val_1 (Xho1) F* | *taactcgagAGGATGAAGATGATGATAATAAC* | |
| | *AP2-g Val (AvrII) R* | *taacctaggAATATTCCTGTTGTTTCCCCCTT* | |
| | Ap2G-CRISP_F | tattTTGCCACAAAAAGGAATAAG | |
| | Ap2G-CRISP_R | aaacCTTATTCCTTTTTGTGGCAA | |
| | Ap2g_3UTRgSeq_F | tattGACATACAGTTCTTATAATG | |
| | Ap2g_3UTRgSeq_R | aaacCATTATAAGAACTGTATGTC | |
| | InFu_AP2g_FKBP_F | CAGGAATATTAGCGGCCGCGGAGTGCAGGTGGAAACCAT | |
| | InFu_AP2g_FKBP_R | GGATGATATTTAACGCGTTTCTTCCGGTTTTAGAAGC | |
| | Ap2g_3UTR_PAM_F | ATATTTTTTTTTATACATTATAAGAACTG | |
| | InFu_Ap2g_3UTR_R | CGAAAAGTGCCACCTGACGTCTATGAAATGTTACAGTTTG | |
| | InFu_AP2g_mScarlet_F | CAGGAATATTAGCGGCCGCatggtgagcaagggcgaggc | |
| | InFu_AP2g_mScarlet_R | GGATGATATTTAACGCGTcttgtacagctcgtccatgc | |
| | Ap2g_P2A_Nano_F | CAACAGGAATATTAGCGGCCGCGCCACAAACTTCTCTCTGCT | |
| | Ap2g_NanoLuc_R | GATGATATTTAACGCGTCGCCAGAATGCGTTCGCACAGCC | |
| *Msrp1 (PF3D7_1335000)* | MSRP1_12 gSeq_F | tattGAAATATTATATGAACAAAG | |
| | MSRP1_12 gSeq_R | aaacCTTTGTTCATATAATATTTC | |
| | MSRP1_HDR_F | GTACCGAGCTCgaattcGAACTTCTTACAAATAGCGA | |
| | MSRP1_HDR_R | TGCTCACCATacgcgtAAGTGTATTTAATAAATCCAC | |
| | MSRP1_3UTR_F | GCTGTACAAGaccggtTGAGATACATATGTATATATGTC | |
| | MSRP1_3UTR_R | CGAAAAGTGCCACCTgacgtcTGCATAATTCAATGAATATTCT | |
| | Msrp1_mNeonGreen_F | TAAATACACTTACGCGTATGGTGAGCAAGGGCGAGGAG | |
| | Msrp1_mNeonGreen_R | TATGTATCTCAACCGGTCTTGTACAGCTCGTCCATGCC | |
| **Genotyping** | | | |
| *Ap2-g* | *AP2-g Val_1 Endo F* | *ATGAGTTACAATAATCATTTTCG* | |
| | *AP2-g Val Endo R* | *TATAAATACATTTAGTTATAGGG* | |
| | TGP_Ap2g_Endo_R1 | AATATCCATGTGTAAATAA | |
| | TGP_Ap2g_Endo_F1 | AAATAATAATATCATTAACC | |
| *Msrp1* | CRI_MSRP1_Endo_F | GAACATCAGGATTACAAGGAG | |
| | CRI_MSRP1_Endo_R | CCCTACTTGAAGGAAAACCAGA | |
| *Gdv1* | GDV1_recoEndo_F | GATTTTGATTATGTGAGAG | |
| | GDV1_3UTR_Endo_R | AAATGGTATATACACCAAGG | |
| *Stk2 or Stk2*-KO | *P263 (P7)* | *GAAATCTATATAATGAGAAACATA3* | |
| | *P264 (P8)* | *ATTCTCTCTGTCTCCACCTTT3* | |
| | *P261 (P5)* | *CTTATACTTTTGAGTAGTTGT* | |
| | *P262 (P6)* | *TCATAGTTCACATTTCTTGTT* | |
| Plasmid's primer for 5'-integration test | HDR_R | CGAAAAGTGCCACCTGACGTC | |
| | P283 | AGGGTTATTGTCTCATGAGCGG | **This work** |
| | *B65_R* | *GTAGACCCCATTGTGAGTAC (1)* | 1 |

| ***Gene expression*** | | | Ref |
|---|---|---|---|
| *Ap2-g* | Ap2-g_F | CCTGTATACCCTTCTTCGAAAGC (2) | |
| | Ap2-g_R | TCAAAAGTGCTCTCCTTTCGTG (2) | |
| *Surfin 13.1* | Surfin_13.1_F | ACCCGAAGTGACAACATCTCC (2) | |
| | Surfin_13.1_R | TCTCCACGAGTTCCAAGTTTT (2) | |
| *Surfin 1.2* | Surfin_1.2_F | TTTTTCCCTCGATCTCCGCG (2) | |
| | Surfin_1.2_R | GGGTTTGGCCGTACTCTACT (2) | |
| *Hda_* | HDA_F | AATAATGTGGGCGTGCAAGC (2) | |
| | HDA_R | TTCCTTGCATACCACCCGTT (2) | |
| *Arom* | AROM_F | CCAAAACGGGCGTAATGAACA (2) | |
| | AROM_R | GGGTGACTATTCCCATGATCGT (2) | |
| *Ep2* | EP2_F | ATGCGAGAAACATCCAGATGA (2) | |
| | EP2_R | TTACAATCCGACCTACAAAGACT (2) | |
| *Pfge7* | Pfge7_F | GGCCAAAATGAGGATGTTGA (2) | |
| | Pfge7_R | CGCTAGTATTGGGTGAAGCA (2) | |
| *Gexp5* | Gexp5_F | GTGGTTGTTTGAGAAGTGGTGA (2) | |
| | Gexp5_R | ACAGAATCCGTTTGAGATGATGA (2) | |
| *Msrp1* | MSRP1_F | TACCAGGTGCCTTATCAAGTG (2) | |
| | MSRP1_R | CTTGGTTGTGATTCGGTTGATG | |
| *Sbp1* | Sbp1_F | GGCATCTGCAACTACCGAAT (2) | |
| | Sbp1_R | GCTTGAAAAACCGTCATCGT | 2 |
| *Arginyl-tRNA synthetase* | PF3D7_1218600-Fw | AGCTAAAGAGATGCATGTTGGTCATT (1) | |
| | PF3D7_1218600-Rv | GAGTACCCCAATCACCTACATGA | 1 |


(1) Usui M, *et al.* Plasmodium falciparum sexual differentiation in malaria patients is associated with host factors and GDV1-dependent genes. *Nat Commun* **10**, 2140 (2019).

(2) Prajapati SK, *et al.* The transcriptome of circulating sexually committed Plasmodium falciparum ring stage parasites forecasts malaria transmission potential. *Nat Commun* **11**, 6159 (2020).
